# Alzheimer’s Aβ assembly binds sodium pump and blocks endothelial NOS activity via ROS-PKC pathway

**DOI:** 10.1101/2020.12.10.419879

**Authors:** Tomoya Sasahara, Kaori Satomura, Mari Tada, Akiyoshi Kakita, Minako Hoshi

## Abstract

Amyloid β-protein (Aβ) may contribute to worsening of Alzheimer’s disease (AD) through vascular dysfunction, but the actual molecular mechanisms remain controversial. Using *ex-vivo* blood vessels and primary endothelial cells derived from human brain microvessels, we revealed that patient-derived Aβ assemblies, termed amylospheroids (ASPD), exist on the microvascular surface in patient brains and inhibit vasorelaxation through binding to the α3 subunit of sodium, potassium-ATPase (NAKα3) on endothelial cells. Interestingly, NAKα3 also serves as the toxic target of ASPD in neurons. ASPD elicit neurodegeneration through calcium overload, while ASPD suppress vasorelaxation by inhibiting nitric oxide (NO) production. ASPD-NAKα3 interaction on cerebrovascular endothelial cells disturbs the NO release by inactivating endothelial NO synthase through mitochondrial reactive oxygen species and protein kinase C. The findings suggest that ASPD may dually contribute to neuronal and vascular pathologies through binding to NAKα3. Thus, blocking the ASPD-NAKα3 interaction may be a useful target for AD therapy.

## Introduction

Alzheimer’s disease (AD) is characterized by progressive loss of neurons, deposition of aggregated forms of amyloid-β proteins (Aβs), and intracellular formation of neurofibrillary tangles (NFTs). In addition to these neuropathological features, 60-90% of the brain in AD patients exhibit vascular changes such as deposition of Aβ at cerebrovascular vessels (called cerebral amyloid angiopathy (CAA)), leading to a reduction of cerebral blood flow (Binnewijzend et al., 2016; O’Brien, Eagger, Syed, Sahakian, & Levy, 1992), dysfunction of the blood-brain barrier (Yamazaki & Kanekiyo, 2017), induction of vascular inflammation (Suo et al., 1998), and disturbance of angiogenesis (Fischer, Siddiqi, & Yusufaly, 1990), which may precede the onset of the neuropathological changes and cognitive symptoms (Govindpani et al., 2019). Recently, the symptomatic overlap between AD and vascular dementia has been focused, and the vascular biomarkers are expected to improve the clinical diagnosis of AD (Jack et al., 2018). Notably, earlier works from the Nun studies have suggested that symptomatic progression of AD related to Aβ deposition, but not to NFTs, appeared to be significantly modified by the presence of cerebrovascular abnormalities in AD (Snowdon et al., 1997). Therefore, a better understanding of the molecular mechanisms underlying Aβ-related cerebrovascular dysfunction in AD should help us to understand how vascular dysfunction contributes to AD progression, and will open up new therapy.

In studies of the mechanisms of neurodegeneration in AD brains, we purified highly neurotoxic ~30-mer assemblies of Aβ (later termed “amylospheroids” (ASPD)) from human AD brains (Hoshi et al., 2003; Noguchi et al., 2009). We proved that ASPD bind directly to the neuronal isoform of α3 subunit of the sodium pump (sodium, potassium-ATPase α3 (NAKα3)) and cause the death of mature neurons by impairing the pump activity (Ohnishi et al., 2015). ASPD levels in patients’ brains correlate well with disease severity (Ohnishi et al., 2015). Furthermore, ASPD and NAKα3 levels appeared to be inversely correlated in affected brain regions (Ohnishi et al., 2015). Interestingly, an ASPD-binding peptide, which mimics the ASPD-binding region in NAKα3, blocked ASPD neurotoxicity (Ohnishi et al., 2015). This result opens a new possibility for knowledge-based design of peptidomimetics that block the aberrant ASPD-NAKα3 interaction and thereby inhibit neurodegeneration in AD. Surprisingly, NAKα3 was also later reported to serve as a toxic target of misfolded protein assemblies, such as α-synucleins, superoxide dismutase 1 (SOD1), and tau, leading to other neurodegenerative diseases such as Parkinson’s disease and amyotrophic lateral sclerosis (ALS) (Ruegsegger et al., 2016; Shrivastava et al., 2015; Shrivastava et al., 2019). This illustrates the value and generality of NAKα3 impairment in neurodegeneration.

Recently, we established a mature neuron-based system that allows us to chronologically follow ASPD formation in mature neurons (Komura et al., 2019). With this system, we found that ASPD accumulate mainly in the trans-Golgi network of excitatory neurons and are secreted through as-yet-unknown mechanisms, leading to the death of adjacent NAKα3-expressing neurons (Komura et al., 2019). This finding led us to explore the possibility that secreted ASPD may reach the blood vessels and contribute to the cerebrovascular changes in AD brains. Here, by using *in-vitro* blood cell cultures and *ex-vivo* blood vessels, we showed that ASPD bind to NAKα3 in endothelial cells, as we had previously found in neurons (Ohnishi et al., 2015), and inhibit the pump function. But, in contrast to mature neurons, the aberrant ASPD-NAKα3 interaction in endothelial cells induces production of reactive oxygen species (ROS) in mitochondria and activates protein kinase C (PKC). This increases the PKC-phosphorylated inactive form of endothelial nitric oxide (eNOS), and decreases nitric oxide (NO) production. This in turn would suppress the relaxation of blood microvessels, and might cause a reduction of cerebral blood flow and other vascular dysfunctions in AD brains. Thus, we show a new possibility that brain Aβ assemblies accelerate worsening AD pathologies by affecting the cerebrovascular systems via interaction with the sodium pump.

## Results

### ASPD are present in cerebrovascular vessels of AD brain

We have established ASPD-tertiary-structure dependent antibodies that are highly specific and do not detect other Aβ oligomers recognized by a pan-Aβ oligomer A11 antibody (see details in Table S1 in (Noguchi et al., 2009)). Using one of ASPD-specific antibodies, rpASD1 (Noguchi et al., 2009), we first examined whether ASPD accumulate in AD cerebrovascular vessels. As we have reported (Noguchi et al., 2009), ASPD were widely distributed around senile plaques and neurons in the frontal cortex of AD patients (Fig. 1A). In addition to such brain parenchymal staining, we also detected ASPD in most microvessels (see “*” in Fig. 1A left). A high-power view showed that ASPD staining is also present at the inner endothelial surface of the microvessels (marked by arrows in Fig. 1A right), where it is colocalize with Aβ_1-42_ and Aβ_1-40_ stainings (compare arrows in Fig. 1A right with that of Fig. 1B or 1C right). The staining result is consistent with our previous Mass analyses showing that ASPD purified from AD patients’ brains are composed of both Aβ_1-40_ and Aβ_1-42_ (Noguchi et al., 2009). These results suggest that ASPD are present not only in brain parenchyma but also in the endothelial cells of the microvessels in AD brains. Accordingly, in this work, we focused on the effect of ASPD on brain endothelial cells.

**Figure 1.**
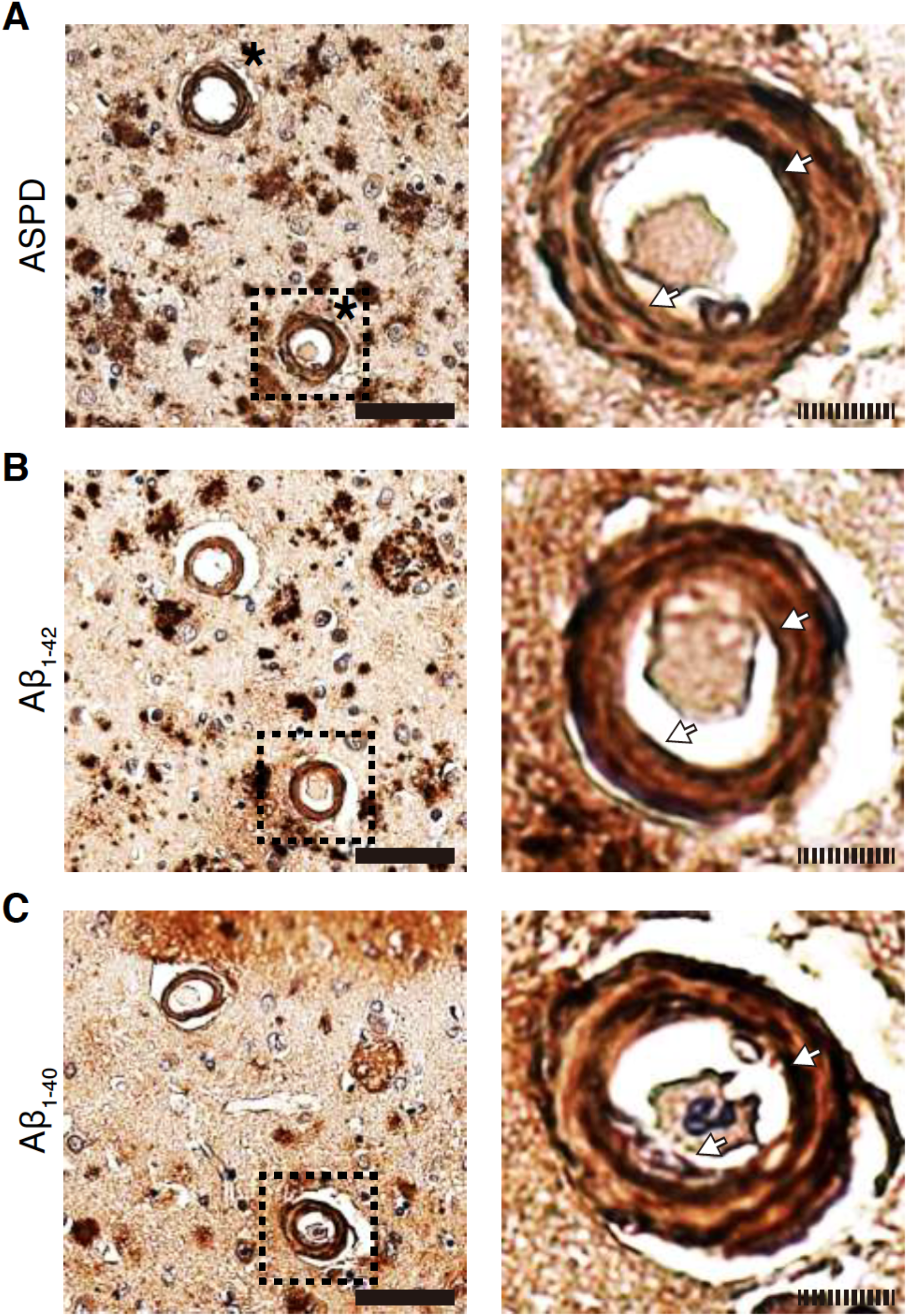
ASPD are present in blood microvessels in AD brains. A-C: Immunohistochemical staining of ASPD (A, ASPD-specific rpASD1 antibody), Aβ_1-4o_ (B), and Aβ_1-42_ (C) in serial sections prepared from frontal cortex of AD patients. Right panels are expanded images of left panels. Scale bars: 50 μm for solid line and 5 μm for hatched line.

### ASPD inhibit relaxation of blood vessels through inhibition of endothelial NAKα3

Endothelial cells release NO, leading to relaxation of blood vessel smooth muscles (Arnold, Mittal, Katsuki, & Murad, 1977; Furchgott & Zawadzki, 1980; Ignarro, Buga, Wood, Byrns, & Chaudhuri, 1987). Therefore, we examined whether ASPD affect the relaxation response of blood vessels. To this end, we used aortic rings isolated from rats, because the aortic rings are considered to be a most sensitive *ex-vivo* vascular model available to detect the change in NO-dependent relaxation response among blood vessels including peripheral arteries and cerebral microvessels (Shimokawa & Godo, 2016). According to the established method (Angus & Wright, 2000; Sasahara, Yayama, Matsuzaki, Tsutsui, & Okamoto, 2013), the isolated aortic rings were contracted by treatment with phenylephrine (an adrenergic α1 receptor agonist), and NO-dependent relaxation was induced by carbachol (a muscarinic M3 receptor agonist). On mature neurons, ASPD affect in a dose-dependent manner, and the effect reaches a plateau at ~40 nM (Ohnishi et al., 2015) (note that ASPD concentrations are shown by using the average mass of ASPD, 128 kDa). We therefore used this plateau concentation of ASPD for the present experiments. Treatment of the isolated aortic rings with 46 nM ASPD for 1 hr inhibited the carbachol-induced relaxation response (Fig. 2A upper panel) and doubled the ED_50_ values of carbachol required for relaxation (Fig. 2A bottom table). We also present the ED_10_ values, because the change of blood vessels in the actual brain takes place in a narrow range, as it is directly linked to the blood pressure. As shown in figure 2A, ASPD also doubled the ED_10_. When ASPD were incubated with ASPD-specific mouse monoclonal mASD3 antibody, which blocks ASPD binding to neurons (Noguchi et al., 2009; Ohnishi et al., 2015), the increase in ED_50_ and ED_10_ was completely abolished (compare black and red circles in Fig. 2A). These results show that ASPD directly suppress the NO-dependent relaxation of the blood vessels, probably through affecting endothelial cells. To eliminate the possibility that ASPD directly affect the smooth muscles of the blood vessels, we confirmed that ASPD did not affect the relaxation response induced by papaverine, which directly relaxes blood vessel smooth muscles in an endothelium-independent manner (Lugnier, Bertrand, & Stoclet, 1972; Martin, Furchgott, Villani, & Jothianandan, 1986). Indeed, 46 nM ASPD did not affect either the papaverine-induced relaxation response of the aortic rings (% maximal relaxation induced by papaverine: 100.9 ± 0.5 and 101.4 ± 0.7% with and without ASPD, respectively; n = 5, *P* = 0.46) or the time to reach the maximal relaxation (3.4 ± 0.1 and 3.6 ± 0.3 min with and without ASPD, respectively; n = 5, *P* = 0.69). These results collectively support that ASPD act on endothelial cells.

**Figure 2.**
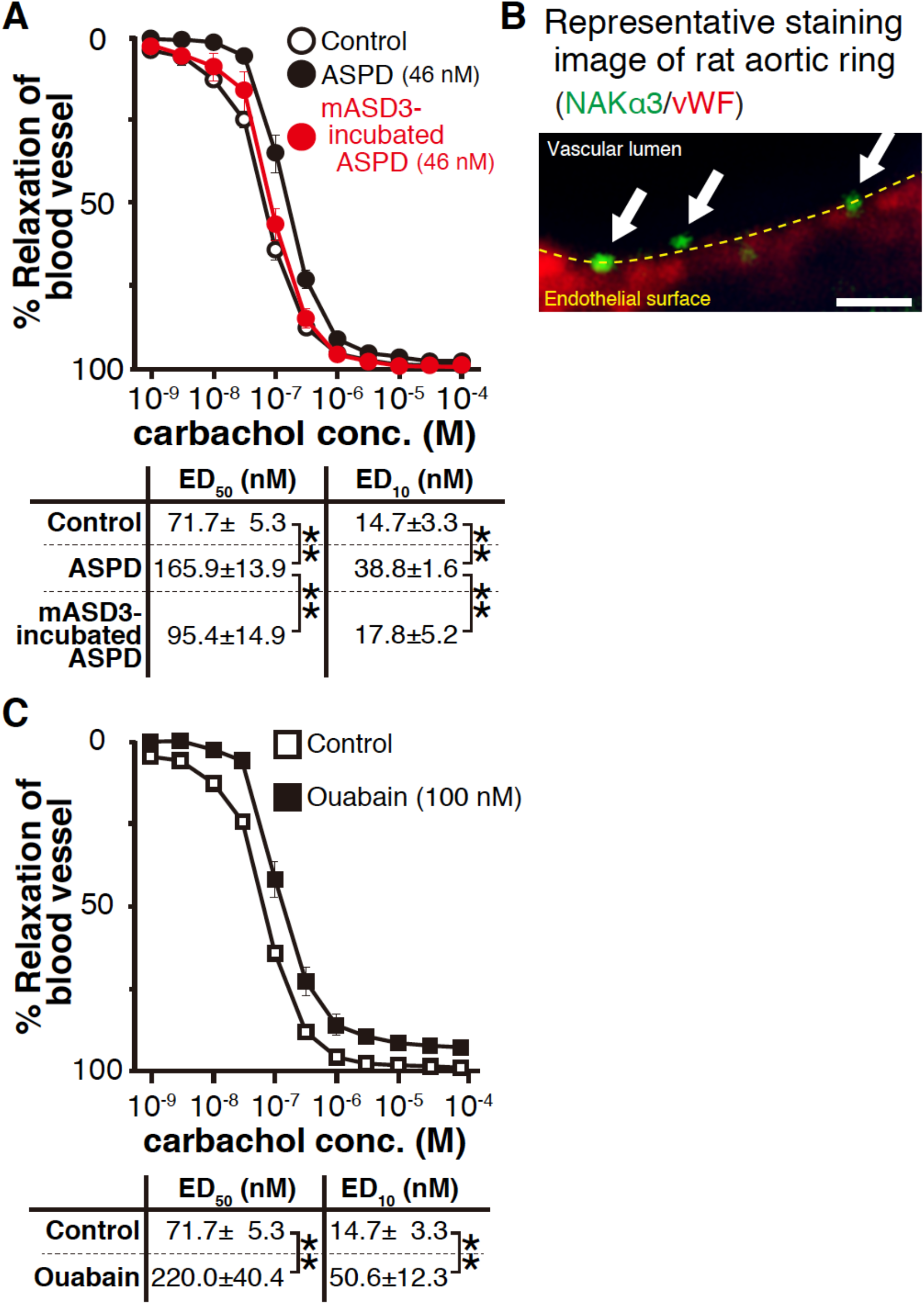
ASPD inhibit *ex-vivo* relaxation response of blood vessels through NAKα3 inhibition. A,C: Effect of 46 nM ASPD (A) or 100 nM ouabain (C) on carbachol-induced relaxation of *ex-vivo* rat aortic rings (n = 3 for mASD3-preincubated ASPD-treated group, and n = 5 for other groups). B: Representative images of immunohistochemical multiple staining of NAKα3 and vWF in the endothelial layer of rat aortic rings. The arrowheads indicate NAKα3 on the apical surface of endothelium. Scale bars: 1 μm. In (A,C), data are presented as means ± S.E. \*\**P* < 0.01 (ANOVA with Scheffé’s method (A) and Welch’s *t*-test (C)).

Next, we elucidated to identify the target protein on endothelial cells to which ASPD bind to inhibit NO release. We speculated that NAKα3 might also serve as an ASPD toxic target in the endothelial cells as it does in mature neurons (Ohnishi et al., 2015). Because NAKα3 is a neuron-specific isoform (Shrivastava, Triller, & Melki, 2018), we first examined whether NAKα3 is present on the endothelial cell surface by immunostaining. We detected patchy NAKα3 staining (green signals indicated by arrows in Fig. 2B) on the vascular lumen surface of the endothelial cells of the isolated aortic rings (red shows a signal of von Willebrand factor (vWF) glycoprotein, an endothelial cytoplasmic marker (Rakocevic et al., 2017)). To confirm the functional involvement of NAKα3 in the suppression of the blood relaxation response, we examined the effect of 100 nM ouabain, a concentration that is enough to inhibit the rodent NAKα3 isoform, but not other rodent isoforms (Noel, Fagoo, & Godfraind, 1990). As shown in figure 2C, this concentration of ouabain sufficiently inhibited the relaxation response and increased both the ED_50_ and the ED_10_ of carbachol required for relaxation of the blood vessels (Fig. 2C), as observed in the ASPD treatment (Fig. 2A). These results collectively support the idea that ASPD suppress blood vessel relaxation by affecting endothelial cell function through inhibition of NAKα3 pump activity. To further dissect the molecular action of ASPD, we decided to use primary cultures of endothelial cells obtained from human brain microvessels.

### ASPD suppress NO production by inactivating eNOS in human primary cerebral endothelial cells

In human brain microvessel-derived endothelial cells, punctate NAKα3 signals were scattered on the cell surface (Fig. 3A; representative 2D image on the left and the vertical views of the z-stack images on the right). Consistently, Western blotting showed that NAKα3 was present in the endothelial cells (green arrowhead in Fig. 3B left). RT-PCR analysis further confirmed NAKα3 expression (green arrowhead in Fig. 3B right). All these results indicated the presence of NAKα3 in human brain microvessel-derived endothelial cells.

**Figure 3.**
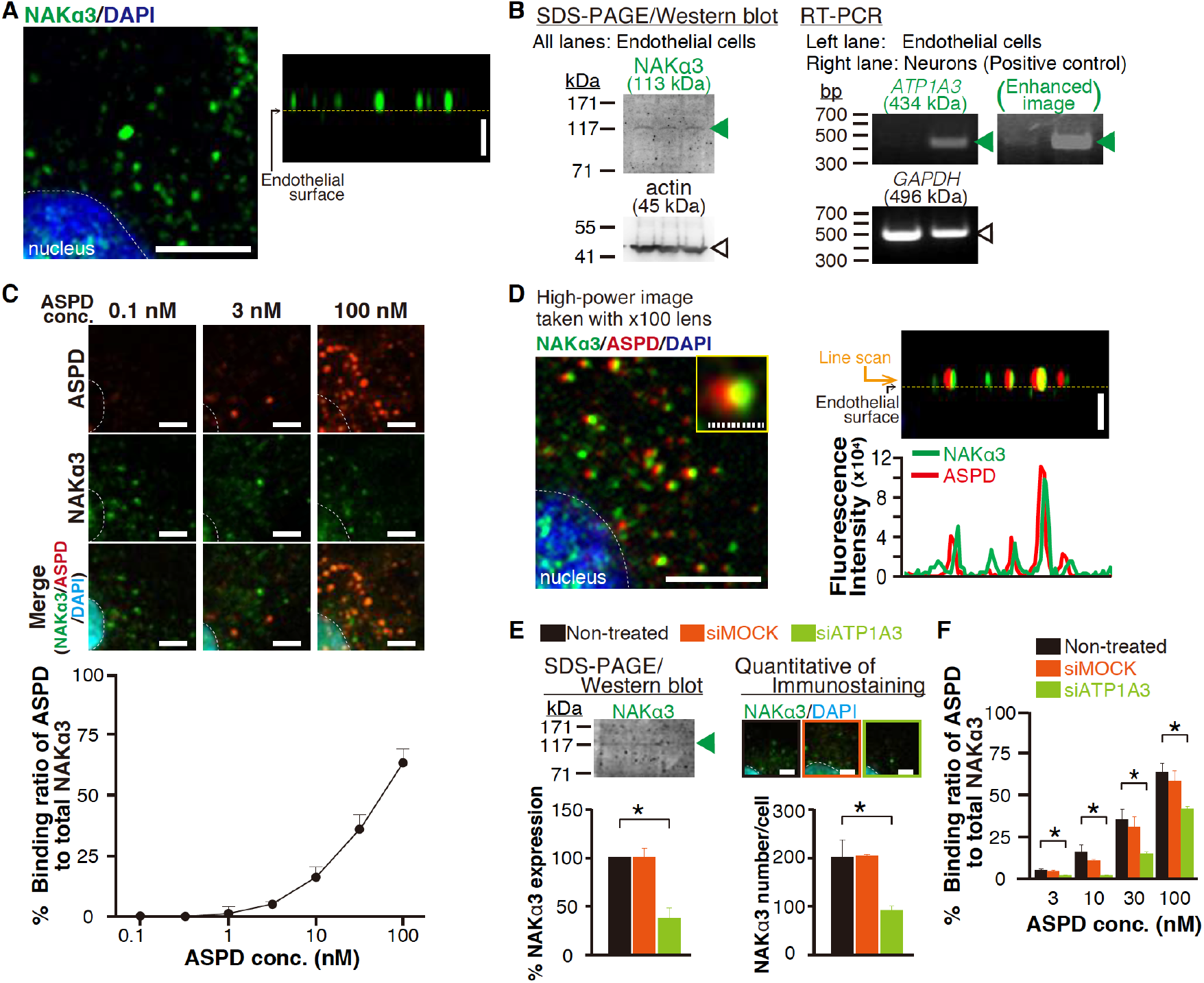
Binding target of ASPD on brain microvessel endothelial cells is NAKα3. A: Representative 2D images of immunocytochemical staining of NAKα3 and nuclei (DAPI) on human primary brain microvessel endothelial cells (left) and vertical section image prepared from z-stack 3D image (right). Scale bars: 5 μm. B: Western blotting for NAKα3 in the endothelial cells (left) and agarose electrophoresis of RT-PCR products for *ATP1A3* mRNA (right). C: Binding ratio of ASPD signal to total NAKα3 signal (the representative images in upper panels and the quantification in bottom, n = 5). Scale bars: 5 μm. D: High-power representative 2D images of multiple immunocytochemical staining of ASPD (ASPD-specific, mASD3 antibody), NAKα3, and nuclei (DAPI) on 30 nM ASPD-treated endothelial cells (left). The vertical section image prepared from z-stack 3D image (right upper) and its line-scan analysis of fluorescence intensities of NAKα3 (green line) and ASPD (red line) (right bottom). Scale bars: 5 μm for solid line and 1 μm for hatched line. E: Western blotting for NAKα3 in the siRNA-transfected endothelial cells (left, n = 3) and quantification of number of punctate NAKα3 signal on the endothelial cells (right, n = 5). Scale bars: 5 μm. F: Effect of siRNA transfection on the binding ratio of ASPD signal to total NAKα3 signal on the endothelial cells (n = 5). The transfection of *ATP1A3* siRNA decreased the ASPD binding ratio to 37 ± 12%, 11 ± 2%, 42 ± 2%, and 66.2 ± 2% in the 3 nM, 10 nM, 30 nM, and 100 nM ASPD-treated groups, respectively. In (C,E,F), data are presented as means ± S.E. \**P* < 0.05 (ANOVA with Scheffé’s method).

We next examined whether ASPD interact with the endothelial NAKα3 using immunostaining. As shown in figure 3C upper panels, binding of ASPD to the endothelial NAKα3 increased dose-dependently (see quantification in Fig. 3C lower panel). Quantification showed that the binding ratio of ASPD to total NAKα3 increased according to the treated ASPD concentrations and reached 63.4 ± 5.7% at 100 nM ASPD (n = 5, Fig. 3C bottom). A high-power image showed that the ASPD and NAKα3 signals are present essentially adjacent to each other, and their centers overlapped well (Fig. 3D left panel and Inset). The vertically sectioned image and its line scan (Fig. 3D right) indicated that the interaction of ASPD and NAKα3 takes place on the endothelial cell surface.

To further confirm ASPD-NAKα3 interaction on the brain endothelial cells, we examined whether knockdown of NAKα3 expression by small interfering RNA (siRNA) blocks the interaction of ASPD and NAKα3. Western blotting and immunostaining consistently showed that the transfection of *ATP1A3* siRNA decreased the amount of NAKα3 to 38 ± 11% (n = 3, *P* = 0.01 compared with siRNA-nontreated group; Fig. 3E left) and to 45 ± 5% (n = 5, *P* = 0.02 compared with siRNA-nontreated group; Fig. 3E right), respectively. In correlation with the decrease of NAKα3 level, the ASPD binding to total NAKα3 was decreased to 31 ± 11% on average (n = 5, Fig. 3F and the representative 2D staining image of 100 nM ASPD-treated cells in Fig. S1). Mock siRNA transfection did not affect either NAKα3 expression (Fig. 3E) or ASPD binding to total NAKα3 (Fig. 3F). The results collectively support that ASPD interact with NAKα3 on the brain endothelial surface.

To elucidate the effect of ASPD on NO production using these human brain endothelial cells, we next determined the ED_50_ of carbachol required for NO release using diaminofluorescein-FM (DAF-FM; Sekisui Medical, Tokyo, Japan), a fluorescent probe for NO quantification (Kojima et al., 1999). As shown in figure S2, a 5-min treatment with carbachol increased NO release dose-dependently, which reached a plateau at around 100 μM. For further experiments, we treated human primary brain microvessel endothelial cells with carbachol at 1 μM, which was approximately the ED_50_ required for NO release (Fig. S2).

As shown in figure 4A, ASPD antagonized carbachol-induced NO release from the human brain microvessel endothelial cells in a dose- and time-dependent manner (Fig. 4A), i.e., the NO release was decreased more rapidly and more strongly in correlation with the increase in ASPD binding ratio to the endothelial NAKα3 (compare Fig. 3C bottom with Fig. 4A). For example, 32 nM ASPD, which interacted with 35 ± 6% of total NAKα3 (n = 4, Fig. 3C bottom), fully inhibited the carbachol-induced NO release after 3 hr incubation (22 ± 11%, n = 4), while 3 nM ASPD, which interacted with 5.2 ± 0.9% of total NAKα3 (n = 4, Fig. 3C bottom), required 6 hr to reach the maximal inhibition (30 ± 18%, n = 4). In contrast, 0.3 nM ASPD, which interacted with 0.7 ± 0.2% of total NAKα3 (n = 4, Fig. 3C bottom), had no effect on the NO release during incubation for up to 6 hr (Fig. 4A). The observed antagonistic effect was attributable to ASPD, as the ASPD-specific mASD3 antibody that inhibits ASPD binding to NAKα3 (Noguchi et al., 2009; Ohnishi et al., 2015) almost completely abolished the effect (Fig. 4B). Binding and functional analyses (Fig. 3 and 4) together support the conclusion that ASPD antagonized carbachol-induced NO release through binding to NAKα3.

**Figure 4.**
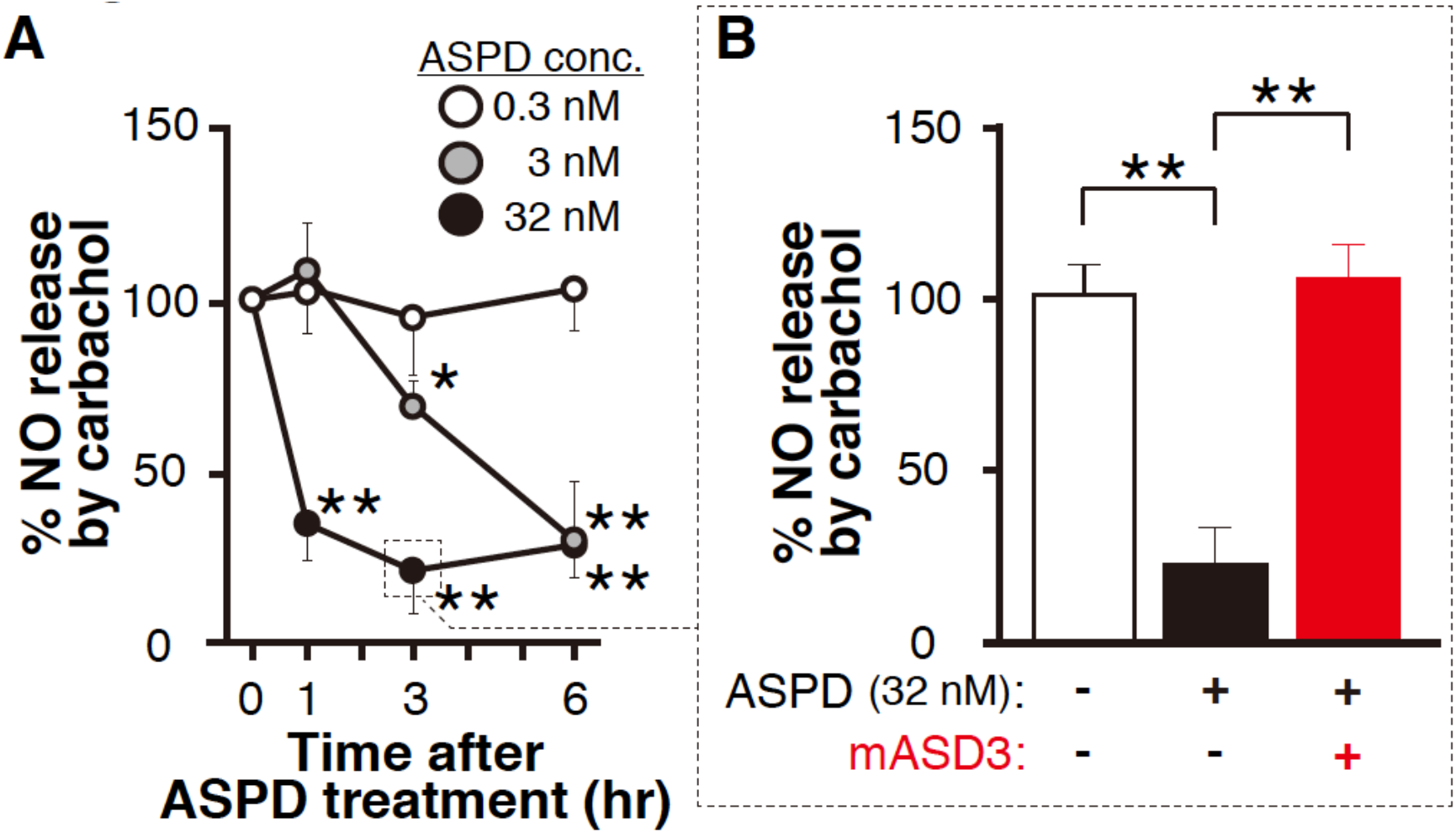
ASPD suppress NO release from brain microvessel endothelial cells. A: Carbachol-induced NO release from human primary brain microvessel endothelial cells treated with ASPD (0.3, 3, or 32 nM for the indicated times) (n = 4). B: Preincubation of ASPD with ASPD-specific mASD3 antibody abolished the 32 nM ASPD-induced suppression of NO release (n = 4). In (A,B), data are presented as means ± S.E. \**P* < 0.05/\*\**P* < 0.01 (ANOVA with Scheffé’s method).

The above findings suggest that ASPD inhibit the activity of eNOS. The eNOS activity is regulated through phosphorylation at Ser^1177^ and Thr^495^ by distinct kinases and phosphatases, respectively (Heiss & Dirsch, 2014); phosphorylation of eNOS-Ser^1177^ (eNOS-P-Ser^1177^) activates, while phosphorylation of eNOS-Thr^495^ (eNOS-P-Thr^495^) deactivates (see scheme in Fig. S3). As for carbachol, it simultaneously activates the kinase responsible for phosphorylating eNOS-Ser^1177^ and the phosphatase responsible for dephosphorylating eNOS-Thr^495^ (see green arrows in Fig. S3). ASPD may antagonize carbachol’s action, as illustrated in figure S3. One possibility is that ASPD disturb the common upstream pathway leading to carbachol-induced activation of the kinase and phosphatase (arrow 1 in Fig. S3). On the other hand, ASPD decrease Ser^1177^ phosphorylation by activating the phosphatase (as shown by arrow 2-1 in Fig. S3) or increase Thr^495^ phosphorylation by activating the kinase (as shown by arrow 2-2 in Fig. S3). Western blotting showed that ASPD, in the absence of carbachol, increased the eNOS-P-Thr^495^ ratio without changing the eNOS-P-Ser^1177^ ratio (Fig. 5A and quantifications below). This suggests that ASPD regulate eNOS-Thr^495^ phosphorylation independently of carbachol (as shown arrow 2-2 in Fig. S3). Consistently, ASPD did not significantly change the eNOS-P-Ser^1177^ ratio even in the presence of carbachol (Fig. 5B). Finally, we confirmed that the ASPD-NAKα3 interaction was responsible for the increase in eNOS-Thr^495^ phosphorylation by knocking down NAKα3 expression with siRNA. As shown in figure 5C, the transfection of endothelial cells with *ATP1A3* siRNA completely blocked the ASPD-induced eNOS-P-Thr^495^ (Fig. 5C). These results collectively show that ASPD-NAKα3 interaction negatively regulates the relaxation response of the blood vessels through eNOS-Thr^495^ phosphorylation independently of the usual relaxation mechanisms induced by carbachol.

**Figure 5.**
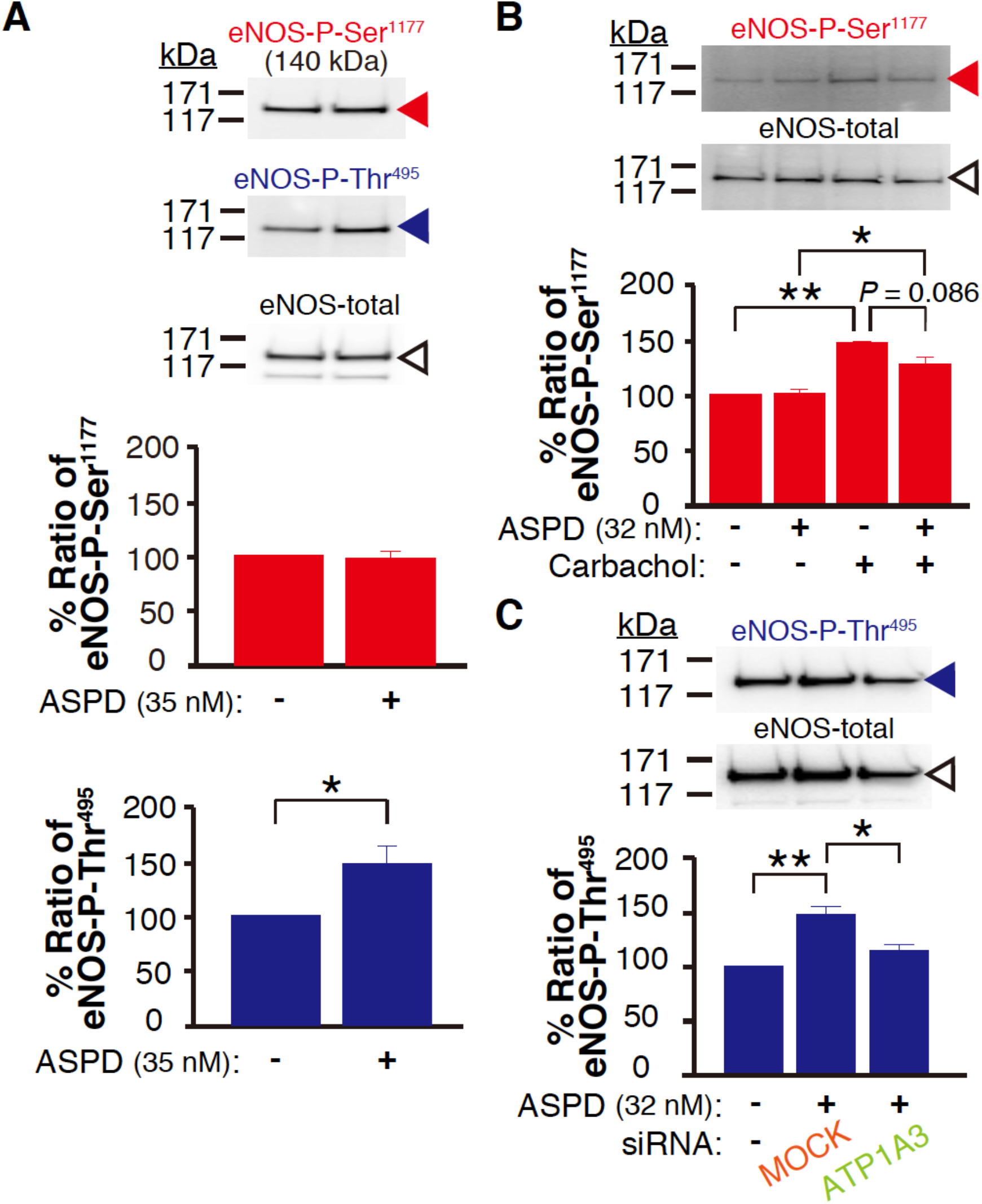
ASPD increase eNOS-P-Ser^1177^ without affecting eNOS-P-Thr^495^. A-C: Western blotting of eNOS-P-Ser^1177^ and eNOS-P-Thr^495^ on 35 nM ASPD-treated human primary brain microvessel endothelial cells (A, n = 4), on 1 μM carbachol-treated endothelial cells after treatment with 32 nM ASPD (B, n = 3), and on 32 nM ASPD-treated endothelial cells transfected with siRNA (C, n = 3). The blotting of NAKα on the endothelial cells with siRNA is shown in Fig. 3E left. In (A-C), data are presented as means ± S.E. \**P* < 0.05/\*\**P* < 0.01 (Welch’s *t*-test (A) and ANOVA with Scheffé’s method (B,C)).

### The mitochondrial ROS/PKC pathway is involved in eNOS-Thr^495^ phosphorylation by ASPD in human cerebral endothelial cells

We next clarified how the ASPD-NAKα3 interaction increases eNOS-Thr^495^ phosphorylation independently of the usual relaxation mechanisms. Previous studies have shown that eNOS-Thr^495^ phosphorylation is mainly regulated by three kinases, protein kinase C (PKC), Rho kinase (ROCK), and AMP-activated protein kinase (AMPK) (Fleming & Busse, 2003; Heiss & Dirsch, 2014). Among the tested inhibitors specific for each kinase, bisindolylmaleimide I (a selective PKC inhibitor) clearly inhibited the ASPD-induced increase in eNOS-P-Thr^495^, but Y-27632 (a ROCK inhibitor) and compound C (an AMPK inhibitor) did not (Fig. 6A). To further confirm the involvement of PKC, we used another inhibitor that works differently: while bisindolylmaleimide I competes at ATP binding site of PKC, calphostin C inhibits the interaction between diacylglycerol and the PKC-regulatory domain (Iida, Kobayashi, Yoshida, & Sano, 1989; Kobayashi, Nakano, Morimoto, & Tamaoki, 1989; Toullec et al., 1991). As shown in figure 6B, calphostin C also completely inhibited the ASPD-induced increase in eNOS-P-Thr^495^. Taken together, these results show that PKC is a major regulator for the ASPD-induced increase in eNOS-Thr^495^ phosphorylation.

**Figure 6.**
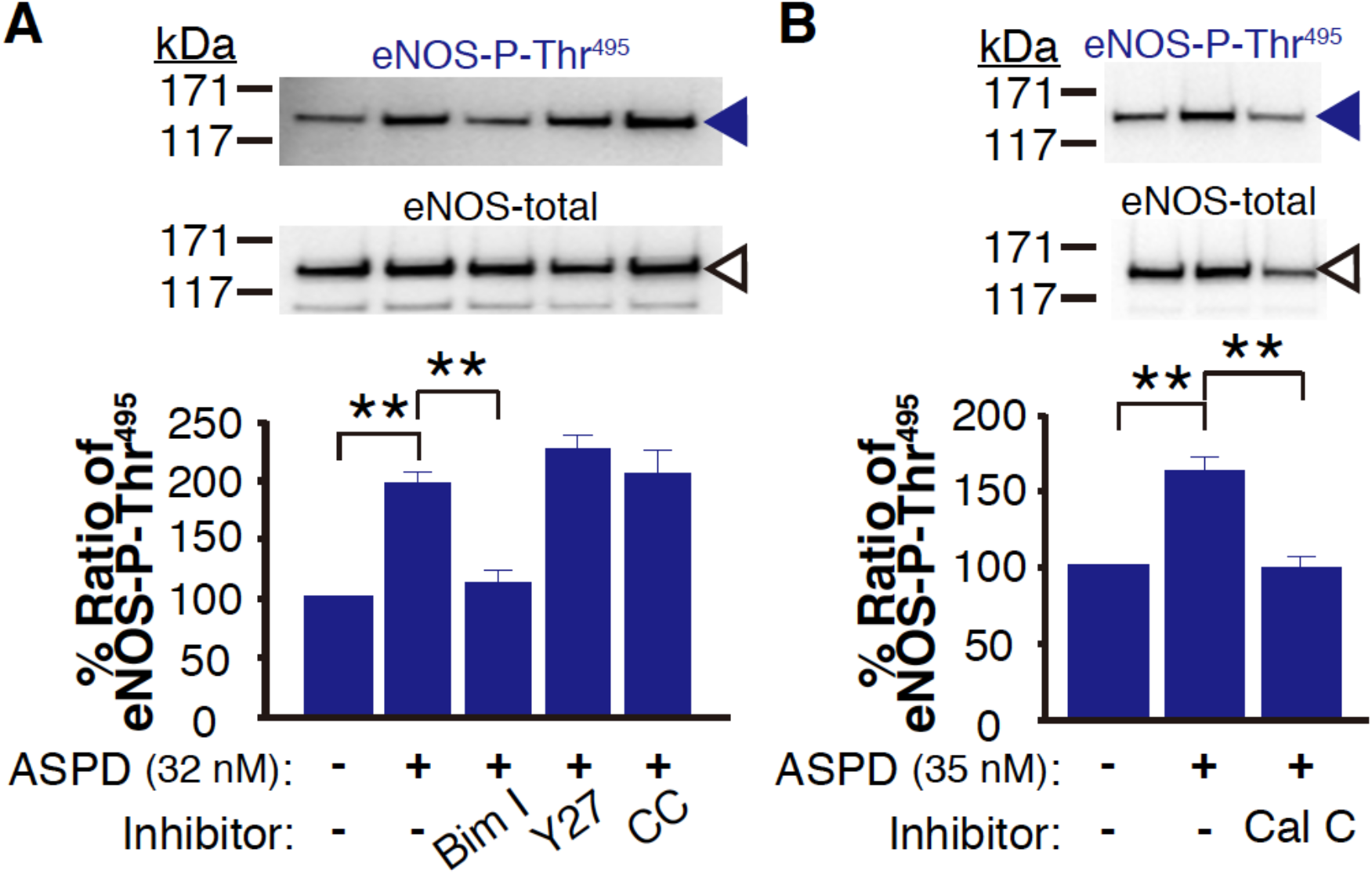
Effects of inhibition of PKC, ROCK, and AMPK on ASPD-induced eNOS-P-Thr^495^. A,B: Western blotting of eNOS-P-Thr^495^ on ASPD (32 nM (A), or 35 nM (B))-treated human primary brain microvessel endothelial cells pretreated with bisindolylmaleimide I (Bim I, 5 μM), Y-27632 (Y27, 10 μM), compound C (CC, 10 μM) (A), or calphostin C. (Cal C, 0.3 μM) (B) (n = 4 (A), and 3 (B)). In (A,B), data are presented as means ± S.E. \*\**P* < 0.01 (ANOVA with Scheffé’s method).

The next question is how the ASPD-NAKα3 interaction leads to PKC activation in the endothelial cells. We elucidated three possible activation mechanisms by using specific inhibitors of each mechanism: tempol (a scavenger of ROS), BAPTA-AM (a chelator of intracellular calcium), or U-73122 (an inhibitor of phospholipase C (PLC)). Because all the tested activation mechanisms are known to be associated with auto-phosphorylation at Ser^660^ in PKC (Cosentino-Gomes, Rocco-Machado, & Meyer-Fernandes, 2012; Feng & Hannun, 1998), the PKC activation was monitored by the ratio of PKC-Ser^660^ phosphorylation (PKC-P-Ser^660^). As shown in figure 7A, only tempol abolished the increase in PKC-P-Ser^660^ associated with PKC activation. Tempol also blocked the increase in eNOS-P-Thr^495^ induced by the ASPD-NAKα3 interaction (Fig. 7B). These results consistently show that the ASPD-NAKα3 interaction induces PKC activation through ROS production.

**Figure 7.**
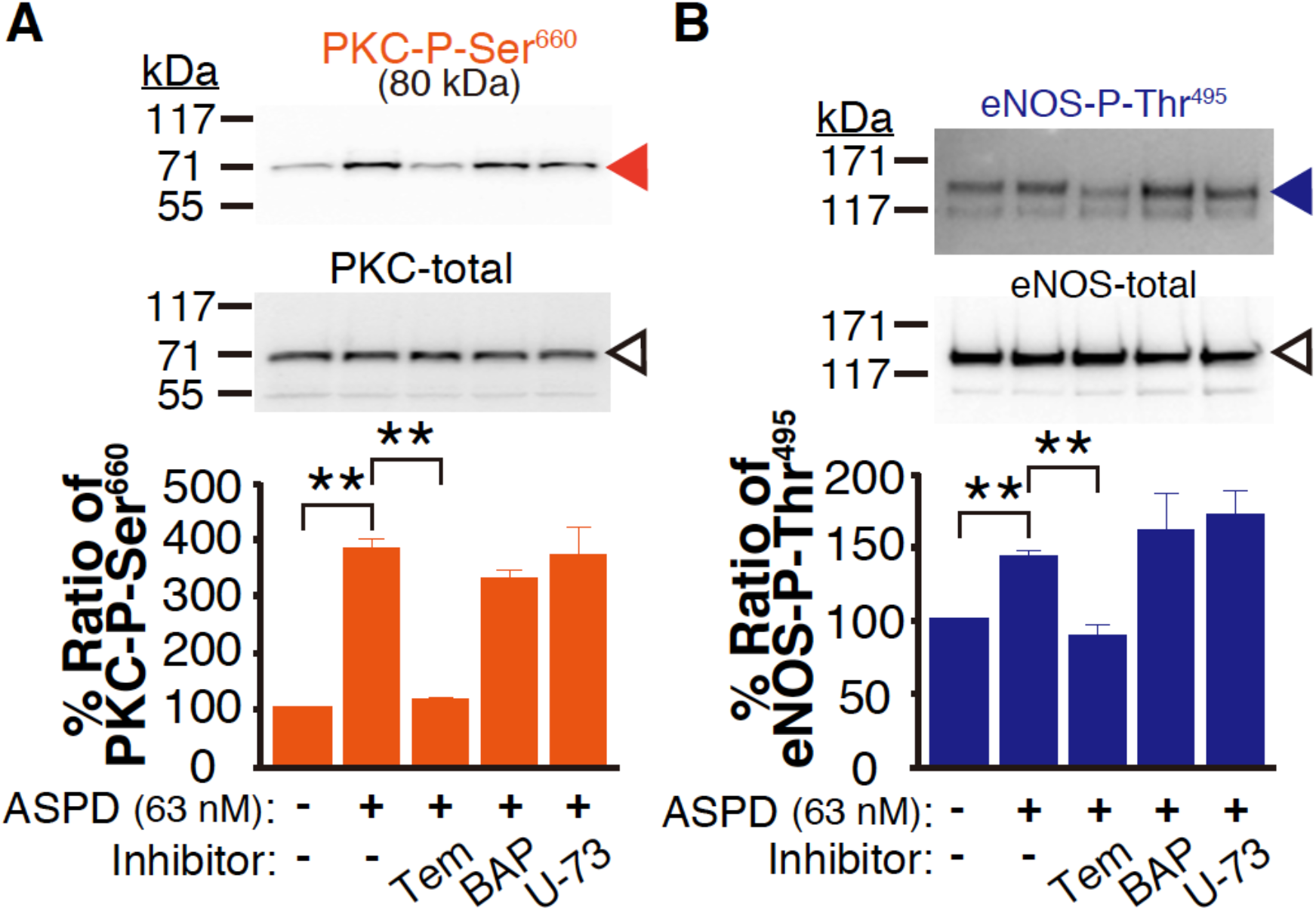
Effects of inhibition of ROS, intracellular calcium, and PLC on the phosphorylation of PKC-Ser^660^ and eNOS-Thr^495^ induced by ASPD. A,B: Western blotting of PKC-P-Ser^660^ (A, n = 4) or eNOS-P-Thr^495^ (B, n = 4) on ASPD (63 nM)-treated human primary brain microvessel endothelial cells pretreated with tempol (Tem, 3 mM), BAPTA-AM (BAP, 30 μM), or U-73122 (U-73, 10 μM). In (A,B), data are presented as means ± S.E. \*\**P* < 0.01 (ANOVA with Scheffé’s method).

The final question is where this ROS production occurs (Santilli, D’Ardes, & Davi, 2015), which we examined by using an ROS indicator, CellROX (Thermo Fisher Scientific, Waltham, MA). As shown in figure 8, CellROX detected an increase in ROS in the cytoplasm of endothelial cells treated with 35 nM ASPD for 6 hr. This increase was wiped out by pretreating the cells with inhibitors of mitochondrial ROS generation, YCG-063 or mito-tempol, but was not affected by pretreatment with NADPH oxidase inhibitors, VAS2870 or apocynin (Fig. 8). Although vascular ROS is also produced by xanthine oxidase (Santilli et al., 2015), the results in figure 8 supported that mitochondria are the major source of the ROS production induced by the ASPD-NAKα3 interaction. The scheme in figure 9 summarizes and illustrates our new findings in this study.

**Figure 8.**
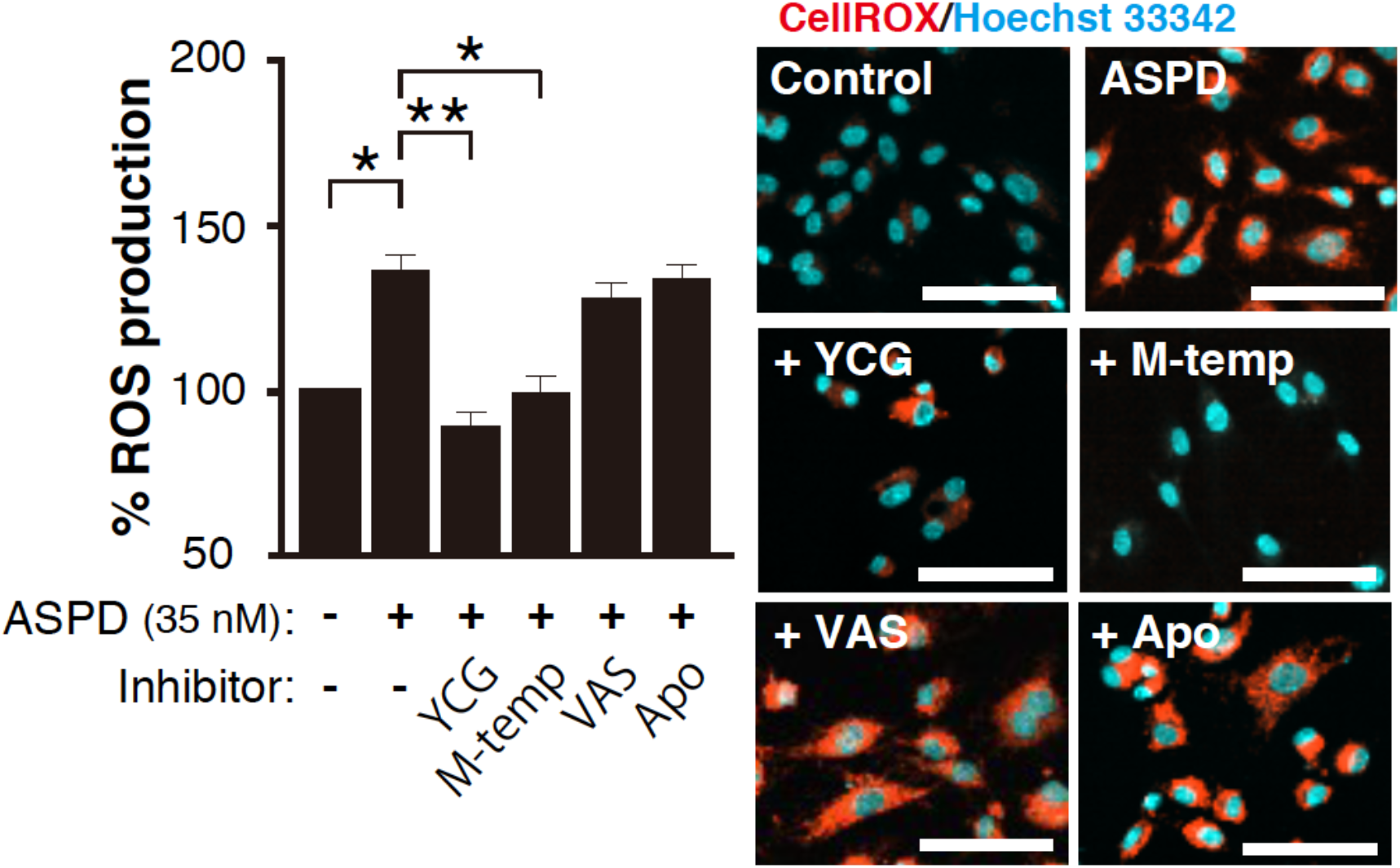
ROS release by ASPD, and the effects of a mitochondrial ROS scavenger and an NADPH oxidase inhibitor. Quantification of ROS release on 35 nM ASPD-treated human primary brain microvessel endothelial cells pretreated with YCG-063 (YCG, 50 μM), mito-tempol (M-temp, 100 μM), VAS2870 (VAS, 10 μM), or apocynin (Apo, 20 μM) (left, n = 4). Representative fluorescence images are shown on the right. Scale bars: 100 μm. Data are presented as means ± S.E. \**P* < 0.05/\*\**P* < 0.01 (ANOVA with Scheffé’s method).

**Figure 9.**
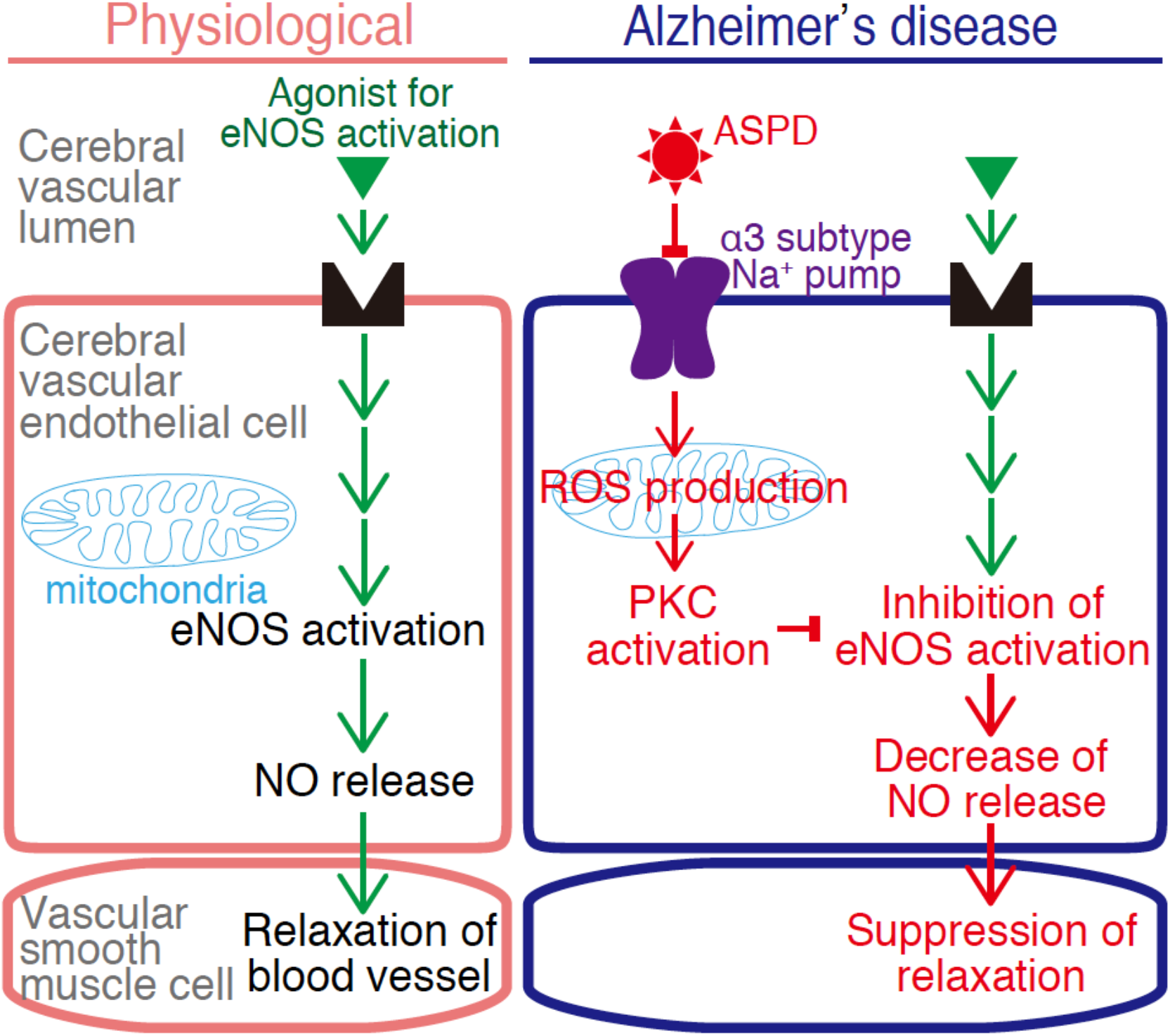
Schematic illustration of the mechanism of ASPD-induced suppression of eNOS activity in brain microvessel endothelial cells. ASPD bind to cell-surface NAKα3 on cerebral microvessel endothelial cells, promote mitochondrial ROS production, activate PKC, increase eNOS-Thr^495^ phosphorylation, and attenuate NO release, resulting in suppression of blood microvessel relaxation response.

## Discussion

A decrease in bioavailable NO leads to cerebrovascular dysfunctions such as reduced cerebral blood flow (Binnewijzend et al., 2016; Boo & Jo, 2003; O’Brien et al., 1992). We found that the aberrant ASPD-NAKα3 interaction in cerebrovascular endothelial cells disturbs NO release by inactivating eNOS through a new system mediated by mitochondrial ROS production, independently of the physiological relaxation system (Fig. 9). The current study opens up a new possibility that ASPD may contribute not only to neurodegeneration in the brain (Komura et al., 2019; Noguchi et al., 2009; Ohnishi et al., 2015), but also to cerebrovascular changes. This is in line with recent observations that vascular changes in the brain contribute to worsening of AD (Govindpani et al., 2019). Such a decrease of cerebral blood flow is responsible for alternating the physiological neurochemical environment, promoting the development of AD-related neuropathology, such as dysfunction and loss of neurons (Zlokovic, 2005). It is noteworthy that ASPD share the same toxic target, NAKα3, in neurons and in cerebral blood microvessels. Therefore, the ASPD-NAKα3 interaction may be a useful target for AD therapy.

The presence of NAKα3 on endothelial cells has not been well investigated, but our immunostaining data revealed the presence of NAKα3 as clusters approximately 230 nm in diameter on the surface of brain microvessel-derived endothelial cells (Fig. 2B and 3A). Moreover, ASPD signals were essentially located proximal to the cellsurface NAKα3 clusters on the endothelial cells (Fig. 3D). This result suggests that NAKα3 may exist in a specific subdomain of the plasma membrane of the brain microvessel-derived endothelial cells, such as membrane microdomains (Leo, Hutzler, Ruddiman, Isakson, & Cortese-Krott, 2020), to which ASPD bind. Western blotting and mRNA analyses also supported the expression of NAKα3 in the cerebrovascular microvessels (Fig. 3B). Notably, ASPD-toxicity-neutralizing antibody and siRNA against NAKα3 clearly abolished the effect of ASPD on the cerebrovascular endothelial cells (Fig. 2A, 3F, 4B, and 5C). Whether NAKα3 is actually present in membrane microdomains or not, and if so, what is its functional role in the physiology of endothelial cells need to await future studies. However, our results collectively demonstrated that ASPD-NAKα3 interaction is responsible for inhibition of eNOS in cerebrovascular endothelial cells.

The treatment of rat endothelial cells with a low concentration of ouabain was reported to increase intracellular calcium (Dong, Komiyama, Nishimura, Masuda, & Takahashi, 2004; Noel et al., 1990). We therefore expected that ASPD-induced PKC activation would occur in an intracellular calcium-dependent manner. However, the intracellular calcium chelator, BAPTA-AM, did not affect PKC activation (Fig. 7A), suggesting that the ASPD-NAKα3 interaction activates PKC through a different pathway. Eventually, we proved that ROS mediates PKC activation by ASPD (Fig. 7A), which was completely abolished by a selective scavenger of mitochondrial ROS production (Fig. 8). The molecular link between NAKα3 inactivation and mitochondrial ROS production remain to be clarified. Interestingly, mitochondrial ROS production was reported to be suppressed by activation of the NAKα3 pump by an NAK-DR region-specific antibody (Yan et al., 2016). These findings together support that NAK activity negatively regulates mitochondrial ROS production.

Suppression of the relaxation response of the blood vessels after Aβ treatment was first reported about two decades ago using rat-derived aortic rings, raising the possibility that Aβ may directly or indirectly reduce NO release (Crawford, Suo, Fang, & Mullan, 1998; Thomas, Thomas, McLendon, Sutton, & Mullan, 1996). Since then, four papers have shown that the eNOS activity is actually decreased after the treatment of blood endothelial cells derived from umbilical vein, aorta, or basilar artery with high concentrations of Aβ (more than 5 μM) (Chisari, Merlo, Sortino, & Salomone, 2010; Gentile et al., 2004; Lamoke et al., 2015; Suhara et al., 2003). These previous studies suggested a possible link between Aβ and eNOS activity regulation, but it has been unknow whether Aβ directly causes the decrease of eNOS activity of the brain microvessels. Moreover, if Aβ does act directly on the eNOS activity of the brain microvessels, there still remained several questions that need to be clarified, such as what molecular entity of Aβ (Aβ monomers or a certain form of Aβ assemblies) actually works, through which target on the blood vessel Aβ acts, and what molecular mechanism lead to the decrease of eNOS activity. Here, we address these questions using human primary endothelial cells derived from the brain microvessels. We previously demonstrated that ~30-mer Aβ assemblies present in AD brains, ASPD (Hoshi et al., 2003; Noguchi et al., 2009), bind to NAKα3 on the endothelial surface, leading to inhibition of eNOS activity by increasing an inactivated state of eNOS phosphorylated at Thr^495^ (Fig. 3, 4, and 5). Thus, by showing that ASPD-specific antibody completely blocked reduction of the eNOS activity observed after ASPD treatment (Fig. 4B), we showed that ASPD directly decreases the eNOS activity by binding to NAKα3. Importantly, while the previous four studies mentioned above observed a decrease in the eNOS-Ser^1177^ phosphorylation after Aβ treatment, which is a part of the physiological pathway for regulating eNOS activity, such a decrease in eNOS-Ser^1177^ phosphorylation was not detected in the case of ASPD (Fig. 5B). Instead, as described above, we found that ASPD activate PKC, which increases eNOS-Thr^495^ phosphorylation, through mitochondrial ROS production (Fig. 6, 7 and 8). PKC has been reported to decrease eNOS-Ser^1177^ phosphorylation by activating Ser/Thr protein phosphatase 2A (PP2A) (Michell et al., 2001). Nevertheless, ASPD appeared not to significantly affect the level of eNOS-Ser^1177^ phosphorylation (Fig. 5B). Thus, ASPD is likely to regulate the contraction of blood vessels in a way different from the physiological pathway for NO release which was previously reported.

To further clarify the molecular link between cerebrovascular dysfunction and parenchymal neuronal damage in the onset of AD induced by Aβ assemblies, one of the essential questions sequestered in future is to understand the source of the Aβ assemblies in the blood microvessels. In the case of ASPD, we have already shown that ASPD are selectively formed in excitatory neurons and secreted (Komura et al., 2019). Therefore, it is natural to consider that ASPD are delivered to the cerebral blood vessels through apolipoprotein E (ApoE), clusterin, or brain meningeal lymphatics, as reported previously (Beeg et al., 2016; Da Mesquita et al., 2018; Garai, Verghese, Baban, Holtzman, & Frieden, 2014; Nelson, Sagare, & Zlokovic, 2017). However, other possibilities, e.g. formation of ASPD in the cerebral blood microvessels, cannot be excluded, because Aβ precursor protein (APP) is also expressed on the surface of cerebral endothelial cells, even though the APP isoform profiles in endothelial cells are different from those in neurons (Kitazume et al., 2010). Even if ASPD are formed in cerebral blood microvessels, as they are in neurons, we consider that a pathological trigger leading to ASPD formation in endothelial cells should be different from that in neurons owing to the differences in APP isoforms and APP processing between neurons and endothelial cells (Grinberg, Korczyn, & Heinsen, 2012; Kakuda et al., 2017; Lasiecka & Winckler, 2011). We believe that further studies to identify the origin of cerebral vascular ASPD will not only deepen our understanding of ASPD themselves, but also help to understand the origin of other Aβ assemblies that accumulate in cerebral blood microvessels.

Interestingly, cerebral vascular ROS was reported to decrease cerebral blood flow through pericyte constriction of cerebral blood vessels (Nortley et al., 2019). Here, we found that ASPD induce mitochondrial ROS production in cerebral endothelial cells, but because released ROS could diffuse to affect nearby cells, it seems plausible that ASPD-induced ROS production in endothelial cells would also affect neighboring pericytes and block the physiological relaxation of cerebral blood vessels.

Decreased cerebral blood flow is an apparent risk factor for AD development (Zlokovic, 2005). Blocking the cerebrovascular toxicity of ASPD is thus expected to be an effective target to prevent worsening of AD. We have previously found an ASPD-binding tetrapeptide that is similar to the ASPD-binding domain on NAKα3, and surprisingly found that the treatment of ASPD with this peptide completely abolished ASPD-induced neuronal apoptosis by blocking the interaction of ASPD with NAKα3 (Ohnishi et al., 2015). Therefore, we expect that this may lead to a new AD therapeutic strategy based on dual attenuation of both the vascular and neuronal toxicities of ASPD.

## Materials and Methods

### Ethics

The Bioethics Committees and the Biosafety Committees of Niigata University, Kyoto University, FBRI at Kobe, and TAO Health Life Pharma approved the experiments using human materials (immunohistochemical study). The Animal Care and Experimentation Committees of Kobe Gakuin University, FBRI at Kobe, and TAO Health Life Pharma approved the experiments using animals (immunohistochemical study and the *ex-vivo* vascular experiments).

### Synthetic ASPD preparation

ASPD were prepared from in-house-prepared Aβ_1-42_ as previously described (Ohnishi et al., 2015). Aβ_1-42_ monomer was aggregated to oligomers by slowly rotating a solution in F12 buffer without L-glutamine and phenol red for 16 hr at 4°C. ASPD were then isolated from the fraction that passed through 0.22-μm filters, but was retained on 100-kDa MWCO filters (Sartorius, Tokyo, Japan). In some experiments, ASPD were preincubated with ASPD-specific mASD3 antibody (0.1 mg/mL) for 2 hr at 4°C before the experiments (Ohnishi et al., 2015).

### Immunohistochemical staining of AD brain and rat aorta

Immunohistochemical staining of autopsied brains from AD patients was performed as previously described (Noguchi et al., 2009). Four μm sections of prefrontal cortex were prepared from the formalin-fixed brains embedded in paraffin wax. The sections for ASPD staining were pretreated with microwave radiation, and those for Aβ_1-40_ or Aβ_1-42_ staining were pretreated with formic acid for 5 min. Then, all sections were pretreated with 0.3% H_2_O_2_-methanol for 60 min followed by incubation with normal goat serum for 30 min at r.t., and further pretreated with a blocking kit (Vector Laboratories, Burlingame, CA). These sections were incubated overnight at 4°C with primary antibodies in the presence of normal goat serum in PBS, followed by incubation for 60 min at r.t. with appropriate biotinylated secondary antibodies. Immunoreactivities were detected by the avidin-biotin-peroxidase complex method using a Vectastain ABC kit (Vector Laboratories). Counterstaining was carried out with Mayer’s hematoxylin. Images were viewed by using a light microscope AX80T (Olympus, Tokyo, Japan) and captured with a digital camera DP70 (Olympus).

Paraformaldehyde (PFA)-fixed aortas were prepared from isoflurane-anesthetized Wistar rats (7-week-old, male; Japan SLC, Shizuoka, Japan) and embedded in paraffin wax. Then 4 μm sections were prepared and pretreated with 10 mM citrate buffer for 30 min at 95°C, followed by incubation with normal goat serum for 30 min at r.t. These sections were incubated overnight at 4°C with primary antibody against NAKα3 (ANP-003, 1:200; Alomone Labs, Jerusalem, Israel) and von Willebrand factor glycoprotein (sc-365712, 1:50; Santa Cruz Biotechnology, Dallas, TX) in the presence of normal goat serum in PBS, and then incubated with the appropriate Alexa Fluor-conjugated secondary antibody (1:1000, Molecular Probes, Waltham, MA) for 1 hr at r.t. Counterstaining was carried out with 4’,6-diamidino-2-phenylindole (DAPI, 1:500; Dojindo Molecular Technologies, Kumamoto, Japan). Fluorescence images were acquired with a confocal laser-scanning microscope LSM710 (Carl Zeiss, Oberkochen, Germany).

### *ex-vivo* relaxation of rat blood vessels

The *ex-vivo* vascular study was performed as previously described with some modifications (Sasahara et al., 2013). Wistar rats (7-week-old, male, Japan SLC) were sacrificed by bleeding from the carotid arteries under isoflurane anesthesia, and the aorta was excised and immediately placed in Krebs-Henseleit solution (118.4 mM NaC1, 4.7 mM KC1, 2.5 mM CaCl2, 1.2 mM KH_2_PO_4_, 1.2 mM MgSO_4_, 25.0 mM NaHCO_3_, and 11.1 mM glucose, 37°C) (K3753; Sigma-Aldrich Japan, Tokyo, Japan). The aorta was cleaned of adherent tissue and cut into 3 mm aortic rings. These rings were vertically fixed under a preload of 1.0 g in a 5 mL organ bath filled with Krebs-Henseleit solution continuously aerated with 95% O_2_/5% CO_2_ gas, and allowed to equilibrate for 60 min. After the pretreatment with ASPD or ouabain for 60 min, the aortic rings were constricted by phenylephrine (1 μM), and then the carbachol (0.001~100 μM)-induced relaxation responses were measured. After the carbachol-induced relaxation, the maximal relaxation responses of the rings evoked by papaverine (0.1 mM) were measured. Relaxation response data are expressed as a percentage of the phenylephrine-induced constriction response. The isometric tension changes of aortic rings were measured with a force-displacement transducer (AP-5; Medical Kishimoto, Kyoto, Japan) coupled to a chart recorder (SS-250 F; SEKONIC, Tokyo, Japan).

### Cell culture

Primary cultures of human endothelial cells derived from brain microvessels (ACBRI 376; Cell Systems, Kirkland, WA) were cultured on collagen I-coated dishes with EGM-2MV medium (CC-3202; Lonza Japan, Tokyo, Japan) at 37°C under 5% CO_2_. The medium of 80% confluent cultures at passage 6 ~ 8 was replaced with fresh medium, and after 24 hr these cultures were employed for the experiments below.

### Immunocytochemical staining of NAKα3 and ASPD

Endothelial cells cultured on collagen I-coated cover glass were treated with ASPD and fixed with 4% (w/v) PFA for 20 min at 37 °C after washing with PBS. The fixed cells were rinsed three times with PBS, treated with 2 mg/ml glycine for 10 min at r.t., rinsed three times with PBS, permeabilized with 0.2% (v/v) Triton X-100 for 5 min at r.t., and rinsed three times with PBS. These cells were pretreated with PBS containing 3% (w/v) BSA (Sigma-Aldrich Japan) and 10% (v/v) normal goat serum (Immuno-Biological Laboratories, Gunma, Japan) for 30 min at r.t., and incubated overnight with primary antibody against ASPD (ASPD-specific mASD3 antibody, 0.1 μg/ml) (Noguchi et al., 2009; Ohnishi et al., 2015) and NAKα3 (sc-16051-R, 0.4 μg/ml, Santa Cruz Biotechnology) at 4 °C. After three washes with PBS, the cells were incubated with the appropriate Alexa Fluor-conjugated secondary antibody (1:1000, Molecular Probes) with counterstaining by DAPI (1:500, Dojindo Molecular Technologies) for 60 min at r.t. The cells were rinsed three times with PBS and mounted with Prolong Gold anti-fade reagent (Invitrogen, Waltham, MA). Fluorescence images were acquired with a confocal laser-scanning microscope LSM710 (Carl Zeiss), and z-stacked images were taken at 2 μm intervals. The quantitative analysis was performed with a confocal quantitative image cytometer CQ1 and CQ1 software (Yokogawa Electric Corp., Tokyo, Japan). Note that the anti-NAKα3 antibody selectively reacts with NAKα3, except for the signals around nucleoli (in the case of non-neuronal cells, including endothelial cells, the anti-NAKα3 antibody shows thick and aggregated signals in nuclei due to non-specific binding (Ohnishi et al., 2015)).

### Western blotting of NAKα3, eNOS, and PKC

The cultures of endothelial cells were treated with ASPD for 6 hr. For assessment of eNOS-Ser^1177^ phosphorylation, some cultures were additionally treated with 1 μM carbachol for 5 min. Then, all cultures were washed twice with D-PBS without Mg^+^ and Ca^2+^ (14249-95; Nacalai tesque, Kyoto, Japan), and the total whole-cell protein was extracted with RIPA buffer (107 mM NaC1, 50 mM Tris-HCl, 5 mM EDTA, 0.1% (w/v) SDS, 0.5% (w/v) sodium deoxycholate, 1% (v/v) NP-40, 1 μg/ml pepstatin, cOmplete Mini (Sigma-Aldrich Japan), and PhosSTOP (Sigma-Aldrich Japan)). SDS-PAGE/Western blotting was performed as previously described with some modifications (Komura et al., 2019). The protein concentration of RIPA lysates was quantified with a BCA Protein Assay Kit (Thermo Fisher Scientific). 10 μg protein/lane was then separated under denaturing conditions on reducing 3~8% Tris-Acetate gels (NuPAGE, Thermo Fisher Scientific), and HiMark protein standard (Thermo Fisher Scientific) was utilized as a protein marker. Bands transferred to 0.2 μm nitrocellulose membranes were blocked with 5% (w/v) skim milk for 1 hr at room temperature, and probed with a primary antibody against eNOS (sc-654, 1:200; Santa Cruz Biotechnology), phosphorylated eNOS (Ser^1177^, 612393, 1:1000; BD Transduction, Franklin Lakes, NJ. Thr^495^, 612707, 1:1000, BD Transduction), PKC (sc-17769, 1:200, Santa Cruz Biotechnology), phosphorylated PKC-Ser^660^ (9371, 1:1000; Cell Signaling Technology Japan, Tokyo, Japan), or actin (MAB1501R, 1:500; Merck-Millipore, Burlington, MA) overnight at 4°C. Bands were detected with SuperSignal West Femto chemiluminescent substrates (Thermo Fisher Scientific) and quantified using a LAS-4000 Mini (GE Healthcare Japan, Tokyo, Japan). To detect NAKα3 (anti-NAK antibody; sc-16052, 1:250, Santa Cruz Biotechnology), we increased the applied protein volume (30 μg protein/lane) and the detection sensitivity by using the combination of biotinylated secondary antibody and avidin-peroxidase. In some experiments, bisindolylmaleimide I (203290, Merck-Millipore), Y-27632 (10005583; Cayman Chemical, Ann Arbor, MI), compound C (171260, Merck-Millipore), calphostin C (208725, Merck-Millipore), tempol (176141, Sigma-Aldrich Japan), BAPTA-AM (196419, Merck-Millipore), or U-73122 (662035, Merck-Millipore) was added to the culture medium 30 min before the ASPD treatment. Quantitative data is shown as densitometric ratios of phosphorylated eNOS (eNOS-P-Ser1177 or eNOS-P-Thr495) to total eNOS, or phosphorylated PKC (PKC-P-Ser^660^) to total PKC.

### RT-PCR of *ATP1A3* mRNA

Total RNA of cultured endothelial cells was extracted using TRIzol reagent (Thermo Fisher Scientific). RNA (0.5 μg) was reverse-transcribed using ReverTra Ace reaction mixture (TOYOBO, Osaka, Japan) with oligo (dT) primer. The reaction mixtures were incubated at 42°C for 20 min, 99°C for 5 min, then 4°C for 5 min to synthesize the first strand of cDNA. The cDNA was then mixed with KOD FX PCR reaction mixture (TOYOBO) with forward and reverse primers for *ATP1A3* (forward primer 5’-CGCCGGGACCTGGATGACCTC-3’ and reverse primer 5’-CGGATCACCAGGGCTTGCTGG-3’ for *ATP1A3*; the PCR product was detected at 434 bp) (Fransen, Hendrickx, Brutsaert, & Sys, 2001) or *GAPDH* (forward primer 5’-CAAGGTCATCCATGACAACTTTG-3’ and reverse primer 5’-GTCCACCACCCTGTTGCTGTAG-3’ for *GAPDH;* the PCR product was detected at 496 bp) (Ouyang et al., 2014_ENREF_42). PCR reaction was performed under the following conditions: initial denaturation at 98°C for 2 min; 30 cycles of denaturation at 98°C for 10 sec, annealing at 55°C for 30 sec, and extension at 68°C for 30 sec; final extension at 68°C for 7 min. PCR products were separated on 1.5% agarose gel and visualized using ethidium bromide.

### Knockdown of NAKα3

*ATP1A3* siRNA (s1724, Thermo Fisher Scientific) was mixed with Lipofectamine 3000 reagent (Thermo Fisher Scientific) according to the manufacturer’s protocol. The mixture was then added to the culture medium on 80% confluent endothelial cultures, and after 6 hr incubation, the culture medium containing siRNA was replaced with fresh EGM-2MV medium. At 72 hr after the siRNA mixture treatment, the endothelial cells were treated with ASPD. Then, the total protein was then extracted with RIPA buffer for SDS-PAGE/Western blot (see above “Western blotting of NAKα3, eNOS, and PKC”), or the endothelial cells were fixed with 4% (w/v) PFA for immunocytochemical staining (see above “Immunocytochemical staining of NAKα3 and ASPD”).

### Measurements of NO release or ROS production

The cultures of endothelial cells treated with ASPD for 1 ~ 6 hr were washed twice with HBSS containing Mg^+^ and Ca^2+^ (09735-75, Nacalai tesque) and then loaded with a fluorescent NO indicator DAF-FM diacetate (SKM423741, Sekisui Medical) or a fluorescent ROS indicator CellROX (Thermo Fisher Scientific), according to the manufacturer’s protocol. NO release was measured in terms of the carbachol-induced enhancement of DAF-FM fluorescence intensity for 5 min detected with CQ1. ROS production was measured in terms of the CellROX fluorescence intensity detected by CQ1. In some experiments, YCG-063 (557354, Merck-Millipore), mito-tempol (18796, Cayman Chemical), VAS2870 (492000, Merck-Millipore), or apocynin (178385, Merck-Millipore) was added into the culture medium at 30 min before the ASPD treatment. The fluorescence data were analyzed using CQ1 software.

### Statistical analyses

All data are expressed as mean ± S.E. We used Statecel2 software (OMS Publication, Tokyo, Japan) for statistical analyses. No data points were excluded from the analysis. Statistical comparisons were performed with the unpaired Welch’s *t*-test between two groups or with one-way analysis of variance (ANOVA) followed by pair-wise comparisons using Scheffé’s method. Differences were considered significant at *P* < 0.05.

## Acknowledgments

We thank Katsutoshi Yayama (Department of Biopharmaceutical Sciences at Kobe Gakuin University, Japan) for help in using the vascular organ chamber; and David B. Teplow (Department of Neurology at University of California, CA) for comments. This work is supported by a Grants-in-Aid for Young Scientists to T. S. (Grant Nos. 6K21713) and by a Grants-in-Aid for Scientific Research to M. H. (Grant Nos. 17H04055) from Japan Society for the Promotion of Science; a Collaborative Research Project of the Brain Research Institute, Niigata University to M.H. (Grant Nos. 2917); and a Life Science Research Grant from Takeda Science Foundation to M. H. (Grant Nos. none).

## Author contributions

T. S. and M. H. designed the research; T. S., K. S., M. T., and M. H. performed the research; all authors analyzed data; T. S. and M. H. wrote the paper.

## Supplemental Figure and corresponding legends

**Figure S1.**
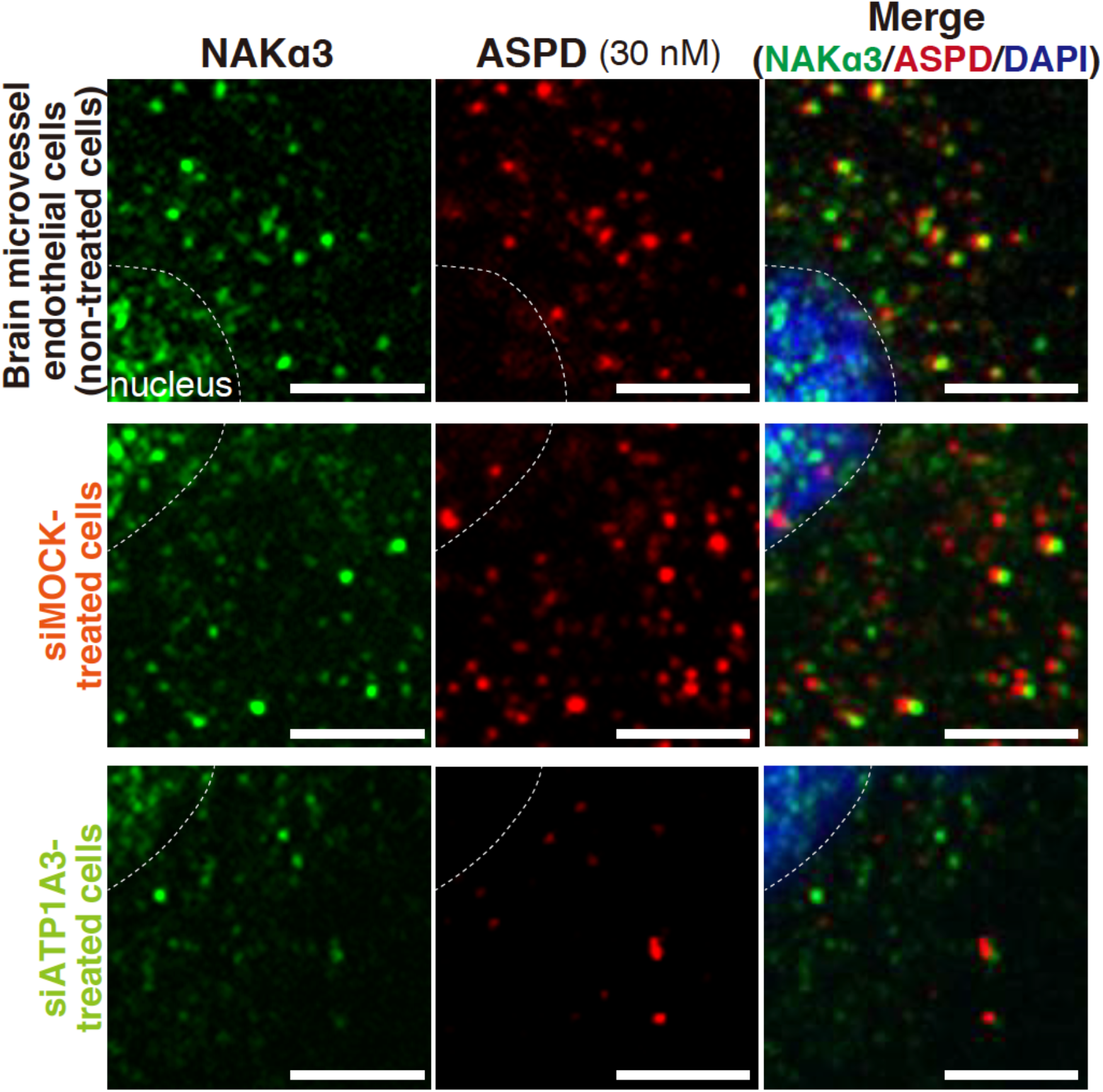
Representative immunocytochemical staining images of 30 nM ASPD-treated brain microvessel endothelial cells in main figure 3F. High-power representative 2D images of immunocytochemical multiple staining of ASPD (ASPD-specific, mASD3 antibody), NAKα3, and nuclei (DAPI) on 30 nM ASPD-treated human primary brain microvessel endothelial cells with or without siRNA transfection (Upper, middle, and bottom panels show non-treated, mock siRNA-treated, and *ATP1A3* siRNA-treated cells, respectively). Scale bars: 5 μm.

**Figure S2.**
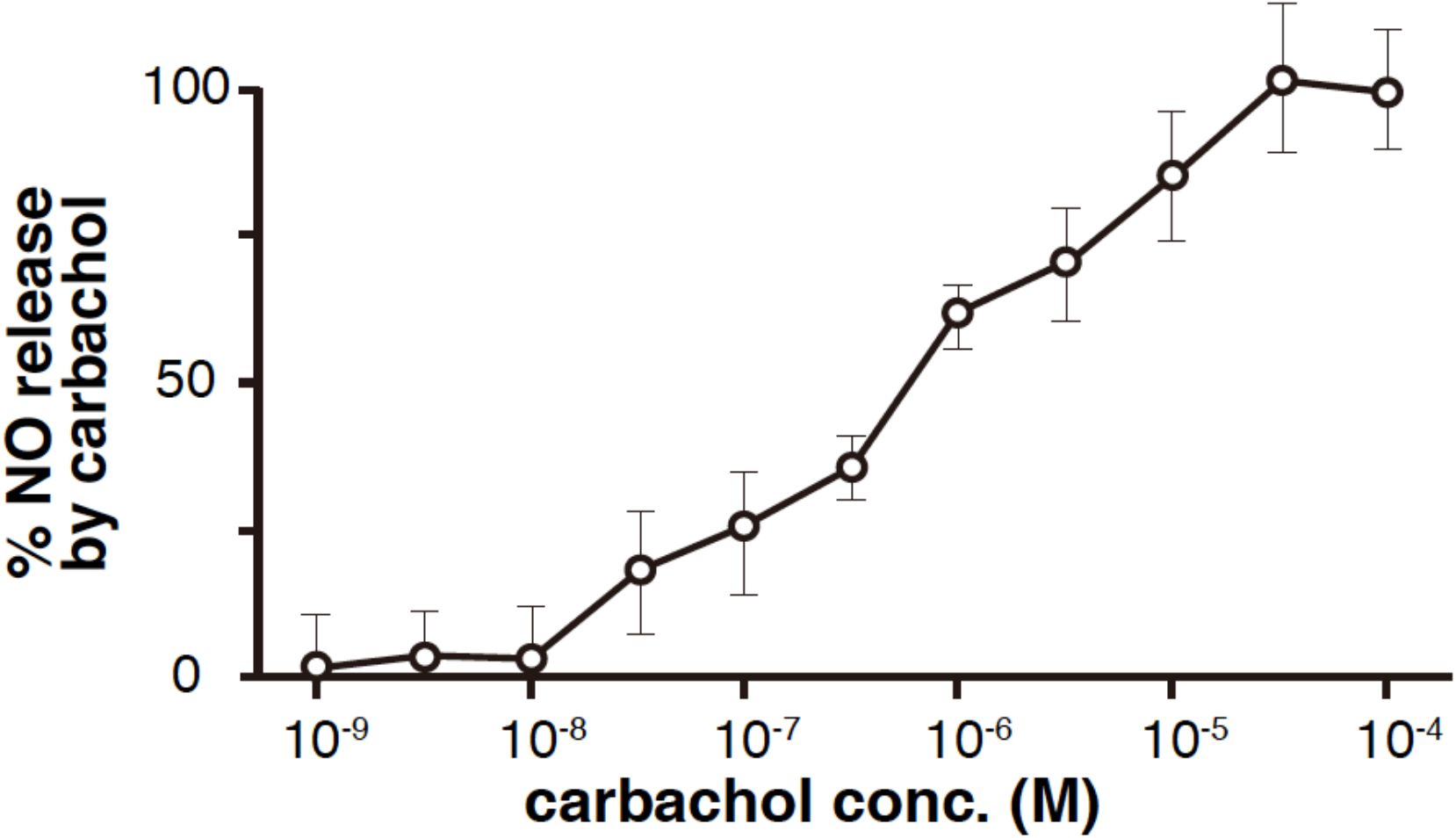
Carbachol concentration-dependent change of NO release from endothelial cells. Change of DAF-FM fluorescent intensity by carbachol was measured (n = 5). A semilog scale was utilized to obtain ED_50_ of carbachol action on NO release. Data are presented as means ± S.E.

**Figure S3.**
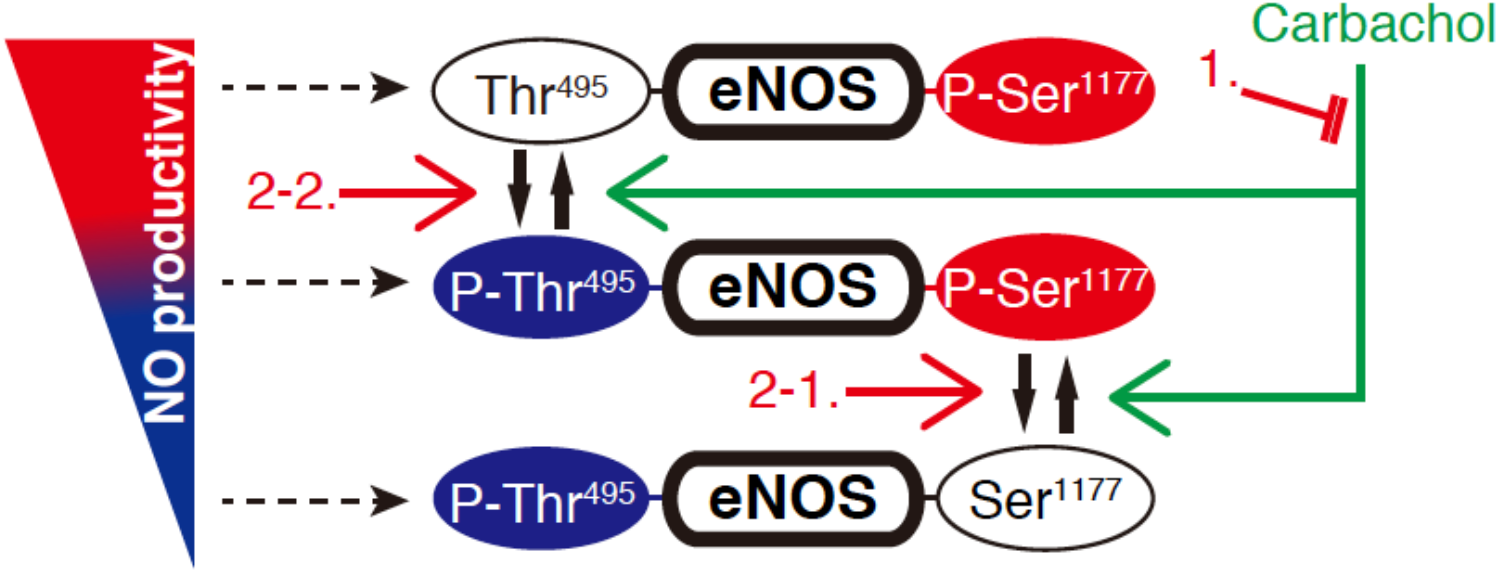
Schematic illustration of the change of eNOS activity. Schematic illustration of the relationship between the NO production by eNOS and the eNOS phosphorylation at Ser^1177^ and at Thr^495^. Carbachol activates eNOS by inducing changes of phosphorylation of both Ser^1177^ and Thr^495^ (green arrows). The three red arrows show possible mechanisms through which ASPD might antagonize carbachol.

## References

Angus, J. A., & Wright, C. E. (2000). Techniques to study the pharmacodynamics of isolated large and small blood vessels. J Pharmacol Toxicol Methods, 44(2), 395–407. doi:10.1016/s1056-8719(00)00121-0

Arnold, W. P., Mittal, C. K., Katsuki, S., & Murad, F. (1977). Nitric oxide activates guanylate cyclase and increases guanosine 3’:5’-cyclic monophosphate levels in various tissue preparations. Proc Natl Acad Sci U S A, 74(8), 3203–3207. doi:10.1073/pnas.74.8.3203

Beeg, M., Stravalaci, M., Romeo, M., Carra, A. D., Cagnotto, A., Rossi, A., … Gobbi, M. (2016). Clusterin Binds to Abeta1-42 Oligomers with High Affinity and Interferes with Peptide Aggregation by Inhibiting Primary and Secondary Nucleation. J Biol Chem, 291(13), 6958–6966. doi:10.1074/jbc.M115.689539

Binnewijzend, M. A., Benedictus, M. R., Kuijer, J. P., van der Flier, W. M., Teunissen, C. E., Prins, N. D., … Barkhof, F. (2016). Cerebral perfusion in the predementia stages of Alzheimer’s disease. Eur Radiol, 26(2), 506–514. doi:10.1007/s00330-015-3834-9

Boo, Y. C., & Jo, H. (2003). Flow-dependent regulation of endothelial nitric oxide synthase: role of protein kinases. Am J Physiol Cell Physiol, 285(3), C499–508. doi:10.1152/ajpcell.00122.2003

Chisari, M., Merlo, S., Sortino, M. A., & Salomone, S. (2010). Long-term incubation with beta-amyloid peptides impairs endothelium-dependent vasodilatation in isolated rat basilar artery. Pharmacol Res, 61(2), 157–161. doi:10.1016/j.phrs.2009.09.004

Cosentino-Gomes, D., Rocco-Machado, N., & Meyer-Fernandes, J. R. (2012). Cell signaling through protein kinase C oxidation and activation. Int J Mol Sci, 13(9), 10697–10721. doi:10.3390/ijms130910697

Crawford, F., Suo, Z., Fang, C., & Mullan, M. (1998). Characteristics of the in vitro vasoactivity of beta-amyloid peptides. Exp Neurol, 150(1), 159–168. doi:10.1006/exnr.1997.6743

Da Mesquita, S., Louveau, A., Vaccari, A., Smirnov, I., Cornelison, R. C., Kingsmore, K. M., … Kipnis, J. (2018). Publisher Correction: Functional aspects of meningeal lymphatics in ageing and Alzheimer’s disease. Nature, 564(7734) E7. doi:10.1038/s41586-018-0689-7

Dong, X. H., Komiyama, Y., Nishimura, N., Masuda, M., & Takahashi, H. (2004). Nanomolar level of ouabain increases intracellular calcium to produce nitric oxide in rat aortic endothelial cells. Clin Exp Pharmacol Physiol, 31(5-6), 276–283. doi:10.1111/j.1440-1681.2004.03995.x

Feng, X., & Hannun, Y. A. (1998). An essential role for autophosphorylation in the dissociation of activated protein kinase C from the plasma membrane. J Biol Chem, 273(41), 26870–26874. doi:10.1074/jbc.273.41.26870

Fischer, V. W., Siddiqi, A., & Yusufaly, Y. (1990). Altered angioarchitecture in selected areas of brains with Alzheimer’s disease. Acta Neuropathol, 79(6), 672–679. doi:10.1007/bf00294246

Fleming, I., & Busse, R. (2003). Molecular mechanisms involved in the regulation of the endothelial nitric oxide synthase. Am J Physiol Regul Integr Comp Physiol, 284(1), R1–12. doi:10.1152/ajpregu.00323.2002

Fransen, P., Hendrickx, J., Brutsaert, D. L., & Sys, S. U. (2001). Distribution and role of Na(+)/K(+) ATPase in endocardial endothelium. Cardiovasc Res, 52(3), 487–499. doi:10.1016/s0008-6363(01)00412-6

Furchgott, R. F., & Zawadzki, J. V. (1980). The obligatory role of endothelial cells in the relaxation of arterial smooth muscle by acetylcholine. Nature, 288(5789), 373–376. doi:10.1038/288373a0

Garai, K., Verghese, P. B., Baban, B., Holtzman, D. M., & Frieden, C. (2014). The binding of apolipoprotein E to oligomers and fibrils of amyloid-beta alters the kinetics of amyloid aggregation. Biochemistry, 53(40), 6323–6331. doi:10.1021/bi5008172

Gentile, M. T., Vecchione, C., Maffei, A., Aretini, A., Marino, G., Poulet, R., … Lembo, G. (2004). Mechanisms of soluble beta-amyloid impairment of endothelial function. J Biol Chem, 279(46), 48135–48142. doi:10.1074/jbc.M407358200

Govindpani, K., McNamara, L. G., Smith, N. R., Vinnakota, C., Waldvogel, H. J., Faull, R. L., & Kwakowsky, A. (2019). Vascular Dysfunction in Alzheimer’s Disease: A Prelude to the Pathological Process or a Consequence of It? J Clin Med, 8(5), 651. doi:10.3390/jcm8050651

Grinberg, L. T., Korczyn, A. D., & Heinsen, H. (2012). Cerebral amyloid angiopathy impact on endothelium. Exp Gerontol, 47(11), 838–842. doi:10.1016/j.exger.2012.08.005

Heiss, E. H., & Dirsch, V. M. (2014). Regulation of eNOS enzyme activity by posttranslational modification. Curr Pharm Des, 20(22), 3503–3513. doi:10.2174/13816128113196660745

Hoshi, M., Sato, M., Matsumoto, S., Noguchi, A., Yasutake, K., Yoshida, N., & Sato, K. (2003). Spherical aggregates of beta-amyloid (amylospheroid) show high neurotoxicity and activate tau protein kinase I/glycogen synthase kinase-3beta. Proc Natl Acad Sci U S A, 100(11), 6370–6375. doi:10.1073/pnas.1237107100

Ignarro, L. J., Buga, G. M., Wood, K. S., Byrns, R. E., & Chaudhuri, G. (1987). Endothelium-derived relaxing factor produced and released from artery and vein is nitric oxide. Proc Natl Acad Sci U S A, 84(24), 9265–9269. doi:10.1073/pnas.84.24.9265

Iida, T., Kobayashi, E., Yoshida, M., & Sano, H. (1989). Calphostins, novel and specific inhibitors of protein kinase C. II. Chemical structures. J Antibiot (Tokyo), 42(10), 1475–1481. doi:10.7164/antibiotics.42.1475

Jack, C. R., Jr., Bennett, D. A., Blennow, K., Carrillo, M. C., Dunn, B., Haeberlein, S. B., … Contributors. (2018). NIA-AA Research Framework: Toward a biological definition of Alzheimer’s disease. Alzheimers Dement, 14(4), 535–562. doi:10.1016/j.jalz.2018.02.018

Kakuda, N., Miyasaka, T., Iwasaki, N., Nirasawa, T., Wada-Kakuda, S., Takahashi-Fujigasaki, J., … Ikegawa, M. (2017). Distinct deposition of amyloid-beta species in brains with Alzheimer’s disease pathology visualized with MALDI imaging mass spectrometry. Acta Neuropathol Commun, 5(1), 73. doi:10.1186/s40478-017-0477-x

Kitazume, S., Tachida, Y., Kato, M., Yamaguchi, Y., Honda, T., Hashimoto, Y., … Taniguchi, N. (2010). Brain endothelial cells produce amyloid {beta} from amyloid precursor protein 770 and preferentially secrete the O-glycosylated form. J Biol Chem, 285(51), 40097–40103. doi:10.1074/jbc.M110.144626

Kobayashi, E., Nakano, H., Morimoto, M., & Tamaoki, T. (1989). Calphostin C (UCN-1028C), a novel microbial compound, is a highly potent and specific inhibitor of protein kinase C. Biochem Biophys Res Commun, 159(2), 548–553. doi:10.1016/0006-291x(89)90028-4

Kojima, H., Urano, Y., Kikuchi, K., Higuchi, T., Hirata, Y., & Nagano, T. (1999). Fluorescent Indicators for Imaging Nitric Oxide Production. Angew Chem Int Ed Engl, 38(21), 3209–3212. doi:10.1002/(sici)1521-3773(19991102)38:21<3209::aid-anie3209>3.0.co;2-6

Komura, H., Kakio, S., Sasahara, T., Arai, Y., Takino, N., Sato, M., … Hoshi, M. (2019). Alzheimer Abeta Assemblies Accumulate in Excitatory Neurons upon Proteasome Inhibition and Kill Nearby NAKalpha3 Neurons by Secretion. iScience, 13, 452–477. doi:10.1016/j.isci.2019.01.018

Lamoke, F., Mazzone, V., Persichini, T., Maraschi, A., Harris, M. B., Venema, R. C., … Mollace, V. (2015). Amyloid beta peptide-induced inhibition of endothelial nitric oxide production involves oxidative stress-mediated constitutive eNOS/HSP90 interaction and disruption of agonist-mediated Akt activation. J Neuroinflammation, 12, 84. doi:10.1186/s12974-015-0304-x

Lasiecka, Z. M., & Winckler, B. (2011). Mechanisms of polarized membrane trafficking in neurons -- focusing in on endosomes. Mol Cell Neurosci, 48(4), 278–287. doi:10.1016/j.mcn.2011.06.013

Leo, F., Hutzler, B., Ruddiman, C. A., Isakson, B. E., & Cortese-Krott, M. M. (2020). Cellular microdomains for nitric oxide signaling in endothelium and red blood cells. Nitric Oxide, 96, 44–53. doi:10.1016/j.niox.2020.01.002

Lugnier, C., Bertrand, Y., & Stoclet, J. C. (1972). Cyclic nucleotide phosphodiesterase inhibition and vascular smooth muscle relaxation. Eur J Pharmacol, 19(1), 134–136. doi:10.1016/0014-2999(72)90090-8

Martin, W., Furchgott, R. F., Villani, G. M., & Jothianandan, D. (1986). Phosphodiesterase inhibitors induce endothelium-dependent relaxation of rat and rabbit aorta by potentiating the effects of spontaneously released endothelium-derived relaxing factor. J Pharmacol Exp Ther, 237(2), 539–547.

Michell, B. J., Chen, Z., Tiganis, T., Stapleton, D., Katsis, F., Power, D. A., … Kemp, B. E. (2001). Coordinated control of endothelial nitric-oxide synthase phosphorylation by protein kinase C and the cAMP-dependent protein kinase. J Biol Chem, 276(21), 17625–17628. doi:10.1074/jbc.C100122200

Nelson, A. R., Sagare, A. P., & Zlokovic, B. V. (2017). Role of clusterin in the brain vascular clearance of amyloid-beta. Proc Natl Acad Sci U S A, 114(33), 8681–8682. doi:10.1073/pnas.1711357114

Noel, F., Fagoo, M., & Godfraind, T. (1990). A comparison of the affinities of rat (Na+ + K+)-ATPase isozymes for cardioactive steroids, role of lactone ring, sugar moiety and KCl concentration. Biochem Pharmacol, 40(12), 2611–2616. doi:10.1016/0006-2952(90)90578-9

Noguchi, A., Matsumura, S., Dezawa, M., Tada, M., Yanazawa, M., Ito, A., … Hoshi, M. (2009). Isolation and characterization of patient-derived, toxic, high mass amyloid beta-protein (Abeta) assembly from Alzheimer disease brains. J Biol Chem, 284(47), 32895–32905. doi:10.1074/jbc.M109.000208

Nortley, R., Korte, N., Izquierdo, P., Hirunpattarasilp, C., Mishra, A., Jaunmuktane, Z., … Attwell, D. (2019). Amyloid beta oligomers constrict human capillaries in Alzheimer’s disease via signaling to pericytes. Science, 365(6450). doi:10.1126/science.aav9518

O’Brien, J. T., Eagger, S., Syed, G. M., Sahakian, B. J., & Levy, R. (1992). A study of regional cerebral blood flow and cognitive performance in Alzheimer’s disease. J Neurol Neurosurg Psychiatry, 55(12), 1182–1187. doi:10.1136/jnnp.55.12.1182

Ohnishi, T., Yanazawa, M., Sasahara, T., Kitamura, Y., Hiroaki, H., Fukazawa, Y., … Hoshi, M. (2015). Na, K-ATPase alpha3 is a death target of Alzheimer patient amyloid-beta assembly. Proc Natl Acad Sci U S A, 112(32), E4465–4474. doi:10.1073/pnas.1421182112

Ouyang, M., Liu, H., Yang, K., Jiang, W., Ding, Q., Yu, X., & Chen, W. (2014). Olecular mechanism underlying the myeloperoxidase induced apoptosis of HUVEC-12 cells. Int J Clin Exp Med, 7(4), 879–885.

Rakocevic, J., Orlic, D., Mitrovic-Ajtic, O., Tomasevic, M., Dobric, M., Zlatic, N., … Labudovic-Borovic, M. (2017). Endothelial cell markers from clinician’s perspective. Exp Mol Pathol, 102(2), 303–313. doi:10.1016/j.yexmp.2017.02.005

Ruegsegger, C., Maharjan, N., Goswami, A., Filezac de L’Etang, A., Weis, J., Troost, D., … Saxena, S. (2016). Aberrant association of misfolded SOD1 with Na(+)/K(+)ATPase-alpha3 impairs its activity and contributes to motor neuron vulnerability in ALS. Acta Neuropathol, 131(3) 427–451. doi:10.1007/s00401-015-1510-4

Santilli, F., D’Ardes, D., & Davi, G. (2015). Oxidative stress in chronic vascular disease: From prediction to prevention. Vascul Pharmacol, 74, 23–37. doi:10.1016/j.vph.2015.09.003

Sasahara, T., Yayama, K., Matsuzaki, T., Tsutsui, M., & Okamoto, H. (2013). Na(+)/H(+) exchanger inhibitor induces vasorelaxation through nitric oxide production in endothelial cells via intracellular acidification-associated Ca2(+) mobilization. VasculPharmacol, 58(4), 319–325. doi:10.1016/j.vph.2012.11.004

Shimokawa, H., & Godo, S. (2016). Diverse Functions of Endothelial NO Synthases System: NO and EDH. J Cardiovasc Pharmacol, 67(5), 361–366. doi:10.1097/FJC.0000000000000348

Shrivastava, A. N., Redeker, V., Fritz, N., Pieri, L., Almeida, L. G., Spolidoro, M., … Triller, A. (2015). alpha-synuclein assemblies sequester neuronal alpha3-Na+/K+-ATPase and impair Na+ gradient. EMBO J, 34(19), 2408–2423. doi:10.15252/embj.201591397

Shrivastava, A. N., Redeker, V., Pieri, L., Bousset, L., Renner, M., Madiona, K., … Melki, R. (2019). Clustering of Tau fibrils impairs the synaptic composition of alpha3-Na(+)/K(+)-ATPase and AMPA receptors. EMBO J, 38(3), e99871. doi:10.15252/embj.201899871

Shrivastava, A. N., Triller, A., & Melki, R. (2018). Cell biology and dynamics of Neuronal Na(+)/K(+)-ATPase in health and diseases. Neuropharmacology, 107461. doi:10.1016/j.neuropharm.2018.12.008

Snowdon, D. A., Greiner, L. H., Mortimer, J. A., Riley, K. P., Greiner, P. A., & Markesbery, W. R. (1997). Brain infarction and the clinical expression of Alzheimer disease. The Nun Study. JAMA, 277(10), 813–817.

Suhara, T., Magrane, J., Rosen, K., Christensen, R., Kim, H. S., Zheng, B., … Querfurth, H. (2003). Abeta42 generation is toxic to endothelial cells and inhibits eNOS function through an Akt/GSK-3beta signaling-dependent mechanism. Neurobiol Aging, 24(3), 437–451. doi:10.1016/s0197-4580(02)00135-5

Suo, Z., Tan, J., Placzek, A., Crawford, F., Fang, C., & Mullan, M. (1998). Alzheimer’s beta-amyloid peptides induce inflammatory cascade in human vascular cells: the roles of cytokines and CD40. Brain Res, 807(1-2), 110–117. doi:10.1016/s0006-8993(98)00780-x

Thomas, T., Thomas, G., McLendon, C., Sutton, T., & Mullan, M. (1996). beta-Amyloid-mediated vasoactivity and vascular endothelial damage. Nature, 380(6570), 168–171. doi:10.1038/380168a0

Toullec, D., Pianetti, P., Coste, H., Bellevergue, P., Grand-Perret, T., Ajakane, M., … et al. (1991). The bisindolylmaleimide GF 109203X is a potent and selective inhibitor of protein kinase C. J Biol Chem, 266(24), 15771–15781.

Yamazaki, Y., & Kanekiyo, T. (2017). Blood-Brain Barrier Dysfunction and the Pathogenesis of Alzheimer’s Disease. Int J Mol Sci, 18(9). doi:10.3390/ijms18091965

Yan, X., Xun, M., Li, J., Wu, L., Dou, X., & Zheng, J. (2016). Activation of Na+/K+-ATPase attenuates high glucose-induced H9c2 cell apoptosis via suppressing ROS accumulation and MAPKs activities by DRm217. Acta Biochim Biophys Sin (Shanghai), 48(10), 883–893. doi:10.1093/abbs/gmw079

Zlokovic, B. V. (2005). Neurovascular mechanisms of Alzheimer’s neurodegeneration. Trends Neurosci, 28(4), 202–208. doi:10.1016/j.tins.2005.02.001

